# Denoising Diffusion MRI: Considerations and implications for analysis

**DOI:** 10.1101/2023.07.24.550348

**Authors:** Jose-Pedro Manzano-Patron, Steen Moeller, Jesper L.R. Andersson, Kamil Ugurbil, Essa Yacoub, Stamatios N. Sotiropoulos

## Abstract

Development of diffusion MRI (dMRI) denoising approaches has experienced considerable growth over the last years. As noise can inherently reduce accuracy and precision in measurements, its effects have been well characterised both in terms of uncertainty increase in dMRI-derived features and in terms of biases caused by the noise floor, the smallest measurable signal given the noise level. However, gaps in our knowledge still exist in objectively characterising dMRI denoising approaches in terms of both of these effects and assessing their efficacy. In this work, we reconsider what a denoising method should and should not do and we accordingly define criteria to characterise the performance. We propose a comprehensive set of evaluations, including i) benefits in improving signal quality and reducing noise variance, ii) gains in reducing biases and the noise floor and improving, iii) preservation of spatial resolution, iv) agreement of denoised data against a gold standard, v) gains in downstream parameter estimation (precision and accuracy), vi) efficacy in enabling noise-prone applications, such as ultra-high-resolution imaging. We further provide newly acquired complex datasets (magnitude and phase) with multiple repeats that sample different SNR regimes to highlight performance differences under different scenarios. Without loss of generality, we subsequently apply a number of exemplar patch-based denoising algorithms to these datasets, including Non-Local Means, Marchenko-Pastur PCA (MPPCA) in the magnitude and complex domain and NORDIC, and compare them with respect to the above criteria and against a gold standard complex average of multiple repeats. We demonstrate that all tested denoising approaches reduce noise-related variance, but not always biases from the elevated noise floor. They all induce a spatial resolution penalty, but its extent can vary depending on the method and the implementation. Some denoising approaches agree with the gold standard more than others and we demonstrate challenges in even defining such a standard. Overall, we show that dMRI denoising performed in the complex domain is advantageous to magnitude domain denoising with respect to all the above criteria.

## 1 INTRODUCTION

Thermal noise is an inherent property in any object populated by electrons, such as the human body or electronic systems, and hence unavoidable in any Magnetic Resonance Imaging (MRI) experiment (Redpath, 1998). Its statistical properties are well characterised in the complex MRI domain (zero-mean Gaussian with equal variance in real and imaginary channels), as well as in the magnitude image domain for single-channel receiver coils (Rician distribution) (Andersen, 1996; Gudbjartsson and Patz, 1995; Henkelman, 1985). However, noise characterisation is less straightforward with modern MRI acquisitions, as it depends on various factors, including the type of reconstruction, filtering, number of receiver coils and type of acceleration used (Dietrich et al., 2008; Sotiropoulos et al., 2013c). In addition to these complexities, noise can create extra challenges for imaging modalities with inherently low signal-to-noise ratios (SNR), such as diffusion MRI (dMRI), where the information of interest relies on the signal attenuation (Jones, 2010). Apart from increasing uncertainty in estimates, an elevated *noise floor* (i.e. the smallest measurable signal given the noise level), can reduce the dynamic range of the dMRI signal and cause biases in measurements (Andersson, 2008; Bernstein, 1989; Jones and Basser, 2004; Laun et al., 2009; Wood and Johnson, 1999). The implications of noise for quantitative modelling of dMRI data become particularly relevant nowadays, given the aspirations towards higher spatial resolutions, higher angular contrasts and shorter scan durations (Moeller et al., 2021b).

For all these reasons, there has been increased interest over recent years in developing denoising approaches for reducing the effects of thermal noise on dMRI datasets (see Suppl. Figure 1). Approaches can be grouped based on whether denoising is applied during acquisition/reconstruction (k-space denoising) versus as a post-reconstruction step (imaging space denoising). The former category of methods relies on modified reconstruction protocols, with Deep Learning (DL)-based reconstruction methods gaining significant interest (see (Pal and Rathi, 2022; Zeng et al., 2021) for reviews). Generally, these are designed to learn an optimal subsampling of the k-space that allows for an accelerated acquisition and considerable reduction of noise introduced in the images. The second category includes post-reconstruction denoising (e.g. (Fadnavis et al., 2020; Manjón et al., 2008; Moeller et al., 2021a; St-Jean et al., 2016; Veraart et al., 2016b) that can be applied to already acquired images without explicit dependencies on specific hardware, bespoke reconstruction and vendor-specific technologies. Post-reconstruction denoising can be defined as any signal processing method that aims to recover the signal components from an already corrupted mixture of signal and noise, thus preserving the useful information and, consequently, increasing the SNR. Among them, several approaches have been developed following the Marchenko-Pastur PCA-based denoising (MPPCA), first introduced in (Veraart et al., 2016b). Extensions that have been proposed adapt this approach to non-Gaussian noise models (Cordero-Grande et al., 2019), usage of complex space data (Moeller et al., 2021a), or extending the use of PCA to non-linear manifolds (Ramos-Llordén, G. et al., 2021) and tensorial data (Olesen, J.L. et al., 2023).

However, the distinction between signal and noise is not trivial and denoising processes generally have to consider a number of trade-offs. For instance, redundancy in the dMRI data over neighbouring samples (both in spatial and angular domains) can be utilised to filter out non-informative components associated with thermal noise (e.g., (Manjón et al., 2013; Moeller et al., 2021a; St-Jean et al., 2016; Veraart et al., 2016b)); however, this patch-based approach can in principle lead to the introduction of covariances and/or resolution loss (Alkinani and El-Sakka, 2017; Fan et al., 2019). In addition, the deviation from Gaussian noise properties (Aja-Fernández et al., 2011; Aja-Fernández and Vegas-Sánchez-Ferrero, 2016) and from noise stationarity in modern MRI acquisitions (Deshmane et al., 2012; Ding, Y. et al., 2010; Griswold et al., 2002; Lustig et al., 2007; Pruessmann et al., 1999), can go against the mathematical assumptions underlying many denoising approaches, such as the Marchenko-Pastur theorem (Veraart et al., 2016a), introducing extra challenges in the implementation and in the estimation of the noise level (Pieciak, T. et al., 2010; Ding, Y. et al., 2010). Predictive modeling approaches (e.g. (Xiang T. et al., 2023; Jurek et al., 2023; Fadnavis et al., 2020; Tian et al., 2022)) can overcome some of these challenges at the expense of more invasive *filtering*, as measurements are directly replaced by deterministic predictions.

Despite progress and interest in this area, gaps in our knowledge still exist in objectively characterising denoising approaches and assessing their efficacy. Previous studies have used a combination of qualitative assessments with a range of quantitative evaluations, which either have been inconsistent throughout the literature, preventing direct comparisons; or have not been comprehensive enough, ignoring important aspects of denoising performance. Given the absence of a gold standard to compare against, a number of approaches have been used before to assess the efficacy of denoising, ranging from qualitative visual assessments of improvements between raw and denoised data and of residual maps to more quantitative gains in SNR to assess the reduction in variance and improvements in precision (Fadnavis et al., 2020, 2022a; Moeller et al., 2021a; St-Jean et al., 2016). The introduction of spatial covariances, i.e., potential resolution loss after denoising, has been considered only in a few studies, in terms of spatial frequency components (Veraart et al., 2016b), or by estimating the loss of contrast between volumes of the same acquisition with same b-values but different gradient orientation in (Moeller et al., 2021a). In addition, previous studies have largely ignored accuracy improvements with respect to noise floor-induced biases and noise-induced signal rectification. Lastly, evaluations throughout the literature have been done using very different datasets, making it difficult to compare methods, and on some occasions, using relatively high-SNR/low-resolution data, which do not necessarily capture indicative performance at low and ultra-low SNR regimes, where denoising algorithms should arguably be more beneficial and necessary.

In this work, we reconsider what a diffusion MRI denoising method *should* and *should not* do and we accordingly define criteria to characterise denoising efficacy for comparing denoising approaches. We provide a comprehensive set of evaluations to characterise performance in different domains, including 1) benefits in raw signal quality and in reducing noise variance, 2) gains in reducing biases and the noise floor, 3) preservation of spatial resolution, 4) convergence at the high-SNR limit, 5) gains in model parameter estimation (precision and accuracy), 6) agreement of denoised data against a gold standard, 7) efficacy in enabling noise-prone applications, such as ultra-high resolution imaging or reduced scan time. To do so, we have acquired (and publicly share) bespoke datasets with multiple repeats that sample different SNR regimes and have complementary signal properties, ranging from standard 2mm isotropic multi-shell data to 0.9mm isotropic data with relatively high *b*-value. We have also defined an in-vivo gold standard using the complex average of multiple repeats, allowing for the first time quantitative comparisons against denoised data. Without loss of generality, we apply a number of previously published denoising algorithms to these datasets, to both magnitude and complex domains, including Non-Local Means (NLM) (Buades et al., 2005), MPPCA in both magnitude (Cordero-Grande et al., 2019; Veraart et al., 2016a) and complex domain, and NORDIC, which operates in the complex domain (Moeller et al., 2021a). We demonstrate that considering all the above criteria comprehensively can provide novel insights into the performance of denoising approaches and their differences. We propose that these aspects are to be considered in both the development and evaluation of new methods, as they highlight aspects with important implications for downstream analysis.

## 2 METHODS

We start by proposing a set of criteria that a principled and well-behaved dMRI denoising algorithm should fulfil. Intuitively, an ideal denoising method should identify and filter as much noise as possible from the raw signal. At the same time, this must be done without perturbing the information-carrying components of the signal and their properties, regardless of the level of noise contained in the measurements. Specifically, we propose that:

1. Denoising should improve raw signal quality by removing noise-related variance.
2. Denoising should preserve the expected statistical distribution properties of the dMRI signal and remove biases introduced by the noise-floor.
3. Denoising should increase the similarity of a dataset against a low-noise gold standard, such as the average of multiple repeats.
4. Denoising should preserve the spatial resolution of the original data.
5. Denoising should provide gains in model parameter estimation, both for precision and accuracy.
6. Denoising should not be detrimental to subsequent processing and performance should converge at the high-SNR limit.
7. Denoising should be transformative in allowing SNR-limited applications that are otherwise infeasible (e.g., at ultra-high spatial resolution).

Having these considerations as a guide, we collected datasets representing different SNR/CNR/spatial resolution trade-offs. We subsequently used them to evaluate the performance of different denoising approaches, as described in the following sections.

### 2.1. Data

All MRI datasets were acquired from scanning the same healthy subject multiple times at the Centre for Magnetic Resonance Research (CMRR), University of Minnesota in a Siemens 3T Prisma MRI System using a 32-channel head coil. Data were obtained following ethical approval from the University of Minnesota Research Ethics Committee and informed consent was provided for all scanning sessions. Diffusion MRI data were acquired at three different spatial resolutions, representing different SNR regimes. Multi-band (MB) acceleration (Moeller et al., 2010) was used in all cases with a SENSE1 reconstruction (Sotiropoulos et al., 2013c). A T1-weighted MPRAGE sequence was acquired at 0.8×0.8×0.8 mm voxel size (TR=2.4 s, TE=2.22 ms) in each scanning session. The dMRI datasets acquired were:

- Dataset A: Six repeats of a 2mm isotropic multi-shell dataset, each implementing the UK Biobank protocol (Miller et al., 2016) (TR=3s, TE=92ms, MB=3, no in-plane acceleration, scan time ∼6 minutes per repeat). For each repeat 116 volumes were acquired: 105 volumes with AP phase encoding direction, such as 5 *b* = 0 *s/mm*^2^ volumes, and 100 diffusion encoding orientations, with 50 *b* = 1000 *s/mm*^2^ and 50 *b* = 2000 *s/mm*^2^ volumes; and 4 *b* = 0 *s/mm*^2^ volumes with reversed phase encoding direction (PA) for susceptibility induced distortion correction (Andersson and Skare, 2002). This dataset represents a relatively medium-to-high SNR regime.
- Dataset B: Five repeats of a 1.5 mm isotropic multi-shell dataset, each implementing an HCP-like protocol in terms of q-space sampling (Sotiropoulos et al., 2013a) (TR=3.23 s, TE=89.2 ms, MB=4 no in-plane acceleration, scan time ∼16 minutes per repeat). For each repeat 300 volumes were acquired: 297 volumes with AP phase encoding direction, such as 27 *b* = 0 *s/mm*^2^ volumes, and 270 diffusion encoding orientations, with 90 *b* = 1000 *s/mm*^2^, 90 *b* = 2000 *s/mm*^2^, and 90 *b* = 3000 *s/mm*^2^ volumes; and 3 *b* = 0 *s/mm*^2^ volumes with PA phase encoding for susceptibility-induced distortion correction. This is a low-to-medium SNR dataset, with relatively high resolution.
- Dataset C: Four repeats of an ultra-high-resolution multi-shell dataset with 0.9mm isotropic resolution (TR=6.569 s, TE=91 ms, MB=3, in-plane GRAPPA=2, scan time ∼22 minutes per repeat). For each repeat 202 volumes were acquired with orientations as in (Harms et al., 2018): 199 volumes with AP phase encoding direction, such as 14 *b* = 0 *s/mm*^2^ volumes, and 185 diffusion encoding orientations, with 93 *b* = 1000 *s/mm*^2^, and 92 *b* = 2000 *s/mm*^2^ volumes; and 3 *b* = 0 *s/mm*^2^ volumes with PA phase encoding for susceptibility-induced distortion correction. This is a very low SNR dataset to represent extremely noisy data that without denoising are expected to be borderline unusable (particularly for the higher *b*).

Each data set was acquired on a different day, to minimise fatigue, but all repeats within a dataset were acquired in the same session. All acquisitions were obtained parallel to the anterior and posterior commissure line, covering the entire cerebrum.

#### 2.1.1. Data processing and denoising

Both magnitude and phase data were retained for each acquisition to allow evaluations of denoising in both magnitude and complex domains. In order to allow distortion correction and processing for complex data and avoid phase incoherence artefacts, the raw complex-valued diffusion data were rotated to the real axis using the phase information. A spatially varying phase-field was estimated and complex vectors were multiplied with the conjugate of the phase. The phase-field was estimated uniquely for each slice and volume by firstly removing the phase variations from k-space sampling and coil sensitivity combination, and secondly by removing an estimate of a smooth residual phase-field. The smooth residual phase-field was estimated using a low-pass filter with a narrowed tapered cosine filter (a Tukey filter with an FWHM of 58%). Hence, the final signal was rotated approximately along the real axis, subject to the smoothness constraints (see further details in the relevant Supplementary section and code available on https://github.com/SPMIC-UoN/EDDEN).

Having the magnitude and complex data for each dataset, denoising was applied using different approaches prior to any pre-processing to minimise potential changes in the statistical properties of the raw data due to interpolations (Veraart et al., 2016b). We used the following four denoising algorithms:

- Denoising in the magnitude domain: i) The Non-Local Means (*NLM*) (Buades et al., 2005) was applied as an exemplar of a simple non-linear filtering method adapted from traditional signal pre-processing. We used the default implementation in DIPY (v.1.7.0) (Garyfallidis et al., 2014), where each dMRI volume is denoised independently. ii) The Marchenko-Pastur PCA (MPPCA) (denoted as |*MPPCA*| throughout the text) (Cordero-Grande et al., 2019; Veraart et al., 2016b), reflecting a commonly used approach that performs PCA over image patches and uses the MP theorem to identify noise components from the eigenspectrum. We used the default MrTrix3 implementation (v.3.4.0) (Tournier et al., 2019).
- Denoising in the complex domain: i) MPPCA applied to complex data (rotated along the real axis), denoted as MPPCA*. We applied the MrTrix3 implementation of the magnitude MPPCA to the complex data rotated to the real axis (we found that this approach was more stable in terms of handling phase images and achieved better denoising, compared to the MrTrix3 complex MPPCA implementation). ii) The NORDIC algorithm (Moeller et al., 2021a), which also relies on the MP theorem, but performs variance spatial normalisation prior to noise component identification and filtering, to ensure noise stationarity assumptions are fulfilled.

Each denoising approach was applied jointly to all volumes of each repeat in each dataset (i.e., not separating b-shells), with the exception of NLM which treats each volume independently. All data, both raw and their four denoised versions, underwent the same pre-processing steps for distortion and motion correction (Sotiropoulos et al., 2013b) using an in-house pipeline (Mohammadi-Nejad et al., 2019) (note that all data described in this work refer to data in image space; the term “raw” is used to differentiate from the denoised versions). To avoid confounds from potential misalignment in the distortion-corrected diffusion native space obtained from each approach, we chose to compute a single susceptibility-induced off-resonance fieldmap using the raw data for each of the Datasets A, B and C; and then use the corresponding fieldmap for all denoising approaches in each dataset so that the reference native space stays the same for each of A, B and C. Note that differences between fieldmaps before and after denoising are small anyway, as the relatively high SNR *b* = 0 *s/mm*^2^ images are used to estimate them. However, these small differences can cause noticeable misalignments between methods and confounds when attempting quantitative comparisons, which we avoid here using our approach. Hence, for each of the Datasets A, B and C, the raw blip-reversed *b* = 0 *s/mm*^2^ were used in FSL’s *topup* to generate a fieldmap (Andersson and Skare, 2002). This was then used into individual runs of FSL’s eddy for each approach (Andersson and Sotiropoulos, 2016) that applied the common fieldmap and performed corrections for eddy current and subject motion in a single interpolation step. FSL’s *eddyqc* (Bastiani et al., 2019) was used to generate quality control (QC) metrics, including SNR and angular CNR for each *b* value. The same T1w image was used within each dataset. A linear transformation estimated using with boundary-based registration (Greve and Fischl, 2009) was obtained from the corrected native diffusion space to the T1w space. The T1w image was skull-stripped and non-linearly registered to the MNI standard space allowing further analysis. Masks of white and grey matter were obtained from the T1w image using FSL’s *FAST* (Jenkinson et al., 2012) and they were aligned to diffusion space.

We performed tractography for some evaluations. After distortion correction, we used a parametric spherical deconvolution approach, considering partial volume, to estimate up to 3 fibre orientations (Jbabdi et al., 2012). The model likelihood assumed Gaussian noise. Subsequently, we used *XTRACT* (Warrington et al., 2020, 2022) to reconstruct 42 major white matter tracts.

### 2.2. Evaluations

Using the above data we performed a number of evaluations for the different denoising algorithms, guided by the criteria we defined before.

#### 2.2.1. Consideration 1: Provide gains on raw signal quality and reduce variance

We used a number of complementary metrics to assess gains in raw signal quality before and after denoising. First, the signal-to-noise ratio (SNR) and angular contrast-to-noise ratio (CNR) were used to evaluate how the average signal/contrast changes against noise variance. The SNR was evaluated using the average and variance of signal across multiple *b* = 0 *s/mm*^2^ volumes. For each *b* value, the angular contrast-to-noise ratio (CNR) was computed using *eddyqc* (Bastiani et al., 2019) that informs of angular contrast across the different diffusion sensitising directions acquired at a given *b* value. Angular CNR was defined as the ratio of the variance of diffusion-weighted signal across all directions at a given *b* value to the variance of the noise. We used the variance of the residuals of the data against a non-parametric Gaussian Process predictor in undistorted space, as implemented in *eddyqc* (Bastiani et al., 2019) to get an estimate of noise variance across a *b* shell. For our experiments, we computed voxel-wise SNR and CNR, and we reported their distributions across white and grey matter.

As complementary to SNR and CNR, we also looked into the power of spherical harmonics before and after denoising, as higher-order components are expected to be more contaminated by noise. We computed the signal attenuation for each dataset, by normalising using the mean *b* = 0 *s/mm*^2^ volume after distortion correction. We then fit real even spherical harmonics (SH) (Descoteaux et al., 2007) up to *L* = 8 order to the signal attenuation for each shell. We computed the power for each order as 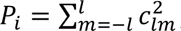, for *l* = 0,2,4,6,8 . Denoising is expected to have a higher influence in reducing the power of higher orders (due to noise removal), compared to lower frequencies.

#### 2.2.2. Consideration 2: Reduce noise-floor and preserve signal statistical properties

Whereas noise in the complex domain is zero-mean Gaussian, dMRI signal produced by modern protocols is noncentral-Chi distributed in the general case (Aja-Fernández et al., 2011; Aja-Fernández and Vegas-Sánchez-Ferrero, 2016). This can lead to an elevated noise floor in the magnitude domain (the minimum measurable signal given the noise level) compared to classical MRI experiments where magnitude signal follows a Rician distribution (Gudbjartsson and Patz, 1995; Salvador et al., 2005). As dMRI information is encoded in the signal attenuation, intensities can be as low as the noise floor in a) regions where the signal attenuation is sufficiently high (e.g. CSF-filled areas or anisotropic regions such as in the Corpus Callosum), and/or b) when b-value and/or spatial resolution are pushed to high limits. In such cases, the signal can be non-linearly rectified leading to biases in a number of extracted features (Jones and Basser, 2004; Sotiropoulos et al., 2013c). The noise-floor can be particularly pronounced in noisy acquisitions, where denoising is expected to be most beneficial. Furthermore, elevated noise floors could be indicative of data that are further away from the Rician distribution regime - typically expected for magnitude MRI data- and are closer to a non-central chi distribution; this can affect any subsequent processing that assumes Gaussian or Rician noise (as a Rician distribution can be typically reduced to a Gaussian for SNR>3 (Salvador et al., 2005).

For all these reasons, it is essential that a denoising approach accounts for and, ideally, mitigates these effects. In order to explore the behaviour of different denoising approaches, we first looked into the distribution of noise intensities before and after denoising. In the case of spatially stationary noise, obtaining such noise distribution from the background voxels of the image would suffice. With multi-channel coils however, noise properties vary spatially (Dietrich et al., 2008). Therefore, we extracted the distribution of noisy signals from the centre of the brain by considering voxels inside the ventricles, where the signal is maximally attenuated and is expected to represent mostly noise. To further explore noise-floor effects in the WM, voxels with very large attenuation, such as in the mid-body of the Corpus Callosum were considered. Due to the large anisotropy in these regions, the signal in these voxels has a large dynamic range (from signal acquired parallel to perpendicular to the fibres) and is also maximally attenuated compared to any other WM region when looking parallel into the main fibre orientation. To better capture and visualise noise floor effects on data acquired with diffusion gradients going from parallel to perpendicular to the main fibre direction we did as follows. We reordered the signal intensities in each voxel in descending order based on the dot product of the corresponding DTI principal eigenvector and the diffusion-sensitising orientations.

#### 2.2.3. Consideration 3: Improve similarity against complex average of multiple repeats

As thermal noise arises from random variations of the signal, powder averaging has been considered the gold-standard for denoising (Moeller et al., 2021a); the noise is averaged out while the signal is preserved. However, this holds only for zero-mean Gaussian noise, i.e. before the magnitude calculation. Otherwise, the noise-floor is preserved after averaging. To avoid this, averaging in the complex domain is needed (Eichner et al., 2015). Having acquired complex data from multiple repeats in each dataset (A, B and C) allows for the calculation of both magnitude-average and complex-average datasets, with the latter being used here as the gold-standard reference for multiple evaluations.

To obtain the averages for each dataset, we first pre-processed the multiple repeats in the magnitude domain. All repeats were concatenated and went through distortion and motion correction using FSL’s *topup* and *eddy*, as described before. This ensured that all volumes across all repeats were aligned together to the first volume of the first repeat using a single interpolation. The aligned and distortion-corrected repeats were then separated and averaged to obtain the magnitude average (denoted as |AVG|). This yielded a dataset where the averaging did not help with the noise floor (i.e., averaging data with a high noise floor will preserve a high noise floor). The distortion fields obtained from the magnitude data were then applied to the corresponding complex data. Aligned and distortion-corrected repeats were then averaged to obtain the complex (real-valued) average (denoted as AVG*). This yielded a dataset where averaging did help with the noise floor (i.e. as if the data had been acquired with higher SNR in the first place). Both magnitude and complex averages were used throughout evaluations to provide references against denoising performance.

#### 2.2.4. Consideration 4: Preserve spatial resolution of the data

Most denoising approaches operate using the principle of redundancy in all sampling domains, where noise at a sample (e.g., voxel) can be determined by using information from other independent samples. Therefore, it is relevant to question whether independent elements in an image change after denoising. As the approaches evaluated make use of spatial patches, we explored whether covariances are introduced after denoising and whether the size of the employed patches can have an effect. We used the idea of identifying smoothness of an image using standardised residuals from a model fit, which is based on Gaussian random field theory and has been commonly used in functional MRI for cluster-based inference (Hayasaka and Nichols, 2003) (Kiebel et al., 1999). By using the spatial covariance of the image partial derivatives computed along a given image dimension, it is possible to identify the degree of smoothness in the data (i.e., effective resolution). More specifically, to estimate resolution, we used FSL’s *smoothest* tool (Jenkinson et al., 2012) (Flitney & Jenkinson 2000) that estimates the smoothness of Gaussian kernels to noise images/model residuals and returns the average Full-Width Half-Maximum (FWHM) as an approximation to the actual spatial resolution. This kernel can be interpreted as the one that would produce the same smoothing as the one we observed in the input images.

We used the residuals after fitting biophysical models to the data, before and after denoising. We fitted two different models to ensure generalisability of findings: the diffusion tensor (DTI) model (Basser and Pierpaoli, 1996) to the lower *b* = 1000 *s/mm*^2^ shell in each case, and the diffusion kurtosis (DKI) model (Jensen et al., 2005) to the *b* = 1000 and *b* = 2000 *s/mm*^2^ shells. We then used the predicted measurements from these models to obtain the residuals and calculate the FWHM of their best Gaussian fit. We repeated this process for the different denoising approaches and at different patch-sizes to explore potential dependencies against patch-size.

#### 2.2.5. Consideration 5: Improve precision and accuracy for model parameter estimation

As denoising should inherently reduce noise variance and noise floor, downstream gains should be expected in dMRI biophysical model estimation for parameter precision and accuracy. We explicitly explored improvements in model parameter accuracy by directly comparing DTI model estimates obtained from the non-denoised and the different denoised datasets, against the respective gold standard ones obtained using the complex average (AVG*). Since we have multiple repeats, we performed the accuracy evaluations across these independent repeats. We also assessed the precision of estimation, by using residual-based bootstrap to calculate the uncertainty of estimated parameters. We specifically used the wild bootstrap (Whitcher et al., 2008) that considers heteroscedasticity in the log-transformed signal, used for ordinary least squares fit of the DTI model, and permutes residuals in-situ. We obtained 250 bootstrap samples and assessed the uncertainty of model parameters across these.

#### 2.2.6. Consideration 6: Converge when denoising at the high-SNR limit

As with any pre-processing approach, denoising algorithms should converge to the limit of high SNR (i.e., where denoising is not really needed), offering larger benefits for low SNR data compared to benefits for high SNR data. Denoising should not be detrimental to any subsequent processing and such convergence provides evidence towards this direction. Data and metrics that converge with and without denoising in the absence of high noise are indicative of a well-conditioned algorithm. Our datasets provide a range of SNR/CNRs, with the 2mm isotropic, multi-shell data (Dataset A) representing what a modern clinical scanner can nowadays provide using bog-standard acquisition choices. This can be considered as mid to high SNR, where denoising is expected to provide fewer gains than for instance bespoke, ultra-high resolution data (Dataset C).

We explored convergence in two ways: a) Using raw signal quality metrics. For each denoising approach, we explored how the CNR gain (before and after denoising) at *b* = 1000 and *b* = 2000 *s/mm*^2^ changed with increasing voxel volume. As our data had higher SNR at low resolution, we expected the CNR gains to decrease with voxel volume. b) Using imaging-derived features. We used data before and after denoising to perform standardised tractography of 42 bundles (Warrington et al., 2020). Using the AVG* as a reference, we then calculated the agreement between the reference tracts and the ones produced after different denoising methods. We did so by computing for each tract the Pearson correlation of the respective path distribution maps (vectorised, after thresholding to voxels with path probabilities higher than 0.5% within a mask given by the same tract estimated with the AVG* data). We calculated this for datasets A and C. We anticipated smaller benefits for the high-SNR Dataset A, compared to lower SNR Dataset C.

#### 2.2.7. Consideration 7: Provide gains for SNR-limited applications

A denoising algorithm should ideally provide gains that are transformative enough in SNR-limited applications. We explored performance in two such scenarios: a) enabling ultra-high spatial resolution imaging or b) reducing scan time by utilising the higher effective SNR per unit time that is expected by denoising (Kawamura et al., 2021).

Firstly, we explored how an approach that relies on consistent quality across the whole brain, like tractography, can operate in the ultra-high resolution (and very noisy) data (Dataset C), before and after denoising. We explored whether we can reconstruct a range of WM bundles, from association, limbic to projection and commissural fibres, that all demonstrate different level of complexity along their route, using the raw and denoised data.

Secondly, we evaluated how denoising can benefit in reducing scan time, by reducing the number of considered diffusion-weighted volumes. We obtained 6 subsets of the original raw data with fewer directions (roughly 15, 35, 50, 65, 80, 100% of the original set of directions) in Datasets B and C. We performed denoising and distortion-correction (motion and eddy current) independently for each of the subsets (using same fieldmap for all of them as before). Subsequently, we obtained DTI metrics (FA and MD) using the *b* = 1000 *s/mm*^2^ volumes, and explored the similarity of the metrics obtained from the subsets (before and after denoising) against the metrics obtained from the gold-standard average of multiple repeats for a particular Dataset.

## 3 RESULTS

In this section, we show results obtained from evaluating the denoising approaches as described above. The following colour code is used throughout: RAW data (non-denoised) in gold, NLM in pink, |MPPCA| (i.e. MPPCA applied to magnitude data) in dark blue, MPPCA* (i.e. MPPCA applied to complex-data) in teal, and NORDIC in green. For some sections, we also show results from the multiple-scans averaged in magnitude (|AVG|) in orange, and multiple-scans averaged in the complex space (AVG*) in red, as baselines to compare against.

### 3.1. Gains in raw signal quality

Qualitative demonstrations of (motion- and distortion-corrected) data before and after denoising for each method and for all the datasets are shown in Figure 1. At higher SNR (Dataset A), differences between raw and denoised data are minimal, mainly noticed for NLM, which seems to provide a smoother image. At lower SNR, a visual improvement in data quality can be perceived after denoising. Noticeably, the signal in the ventricles (which is maximally attenuated and reaches the noise-floor) is different for methods denoising in the magnitude and complex domain space (see arrows in Fig.1). This is particularly evident for the *b* = 2000 s/mm^2^ of the 0.9mm data, where methods that perform denoising in the complex domain (MPPCA*, NORDIC) seem to reduce this signal considerably more compared to the magnitude-domain approaches (NLM, |MPPCA|). Even if this elevated noise floor can be more easily appreciated in the ventricles, it seems to be present in other areas as well, when comparing magnitude vs complex approaches. The same trends are observed for the averages in the complex (AVG*) vs in the magnitude (|AVG|) domain.

**Figure 1.**
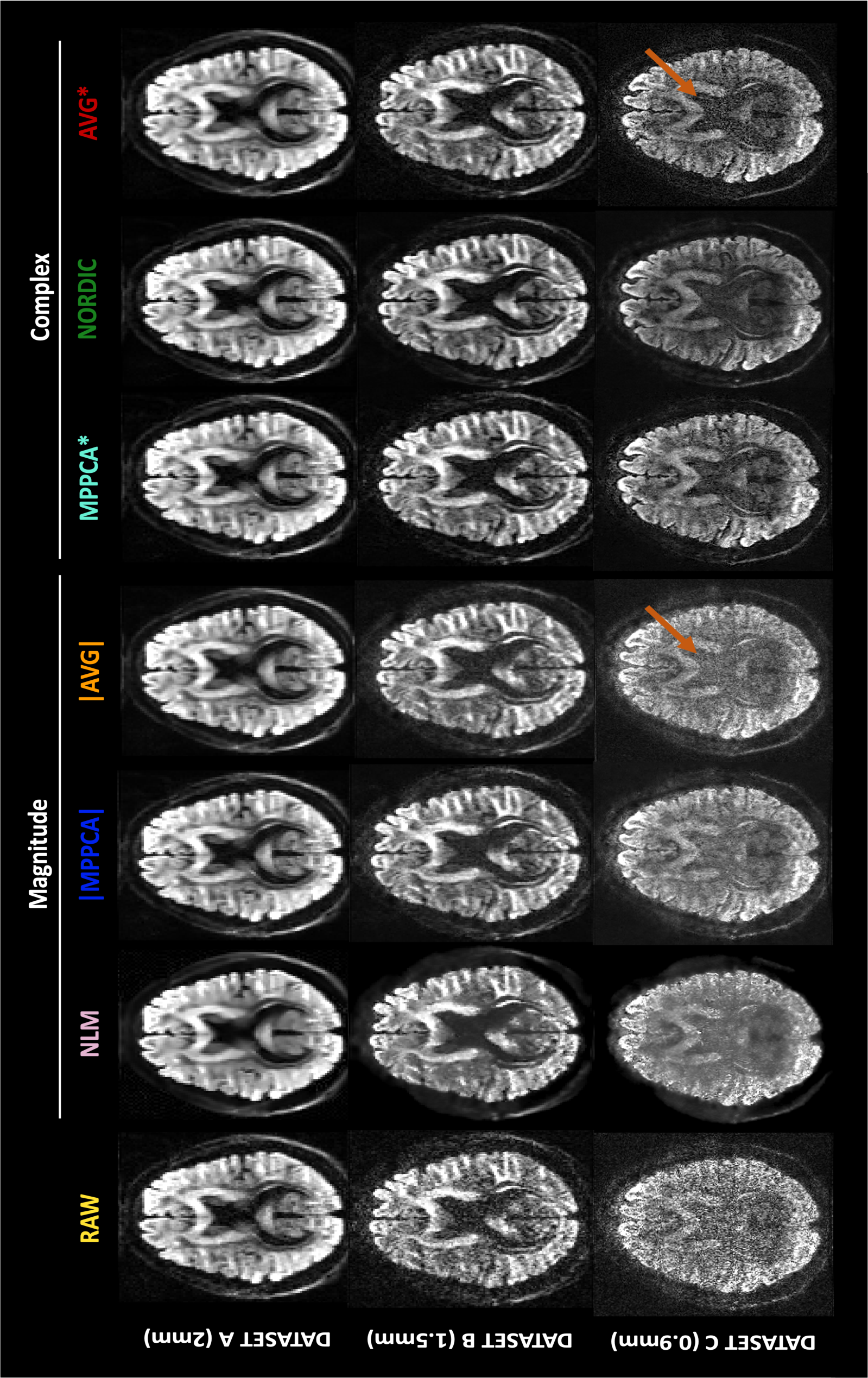
Qualitative maps of the dMRI signals before and after denoising with each method (columns). First row: Dataset A (2mm) in a b = 2000 s/mm^2^ volume. Second row: Dataset B (1.5mm) in a b = 3000 s/mm^2^ volume. Third row: Dataset C (0.9mm) in a b = 2000 s/mm^2^ volume. Arrows point to maximally attenuated signals in the ventricles where the noise-floor can be appreciated.

Gains in raw signal quality are shown in (Figure 2), where boxplots correspond to distributions in the whole brain (white matter distributions are shown in Suppl. Figure 2). Firstly, the results for the raw (i.e. non-denoised) data confirm our anticipation that Dataset A, which is the lowest resolution, had the highest SNR (around 30 for b=0), followed by Dataset B (SNR=15 for b=0 on average) and then the ultra-high resolution Dataset C (SNR=5 for b=0). As expected, all denoising approaches improved SNR and angular CNR for all datasets. NLM showed the largest improvement for b=0 s/mm^2^ volumes; however, this behaviour was not mirrored in denoising DWI volumes. The PCA-based approaches provided the most consistent pattern of improvement across datasets and b-shells, with NORDIC providing the largest gains in CNR, especially at higher b-shells and higher resolutions.

**Figure 2.**
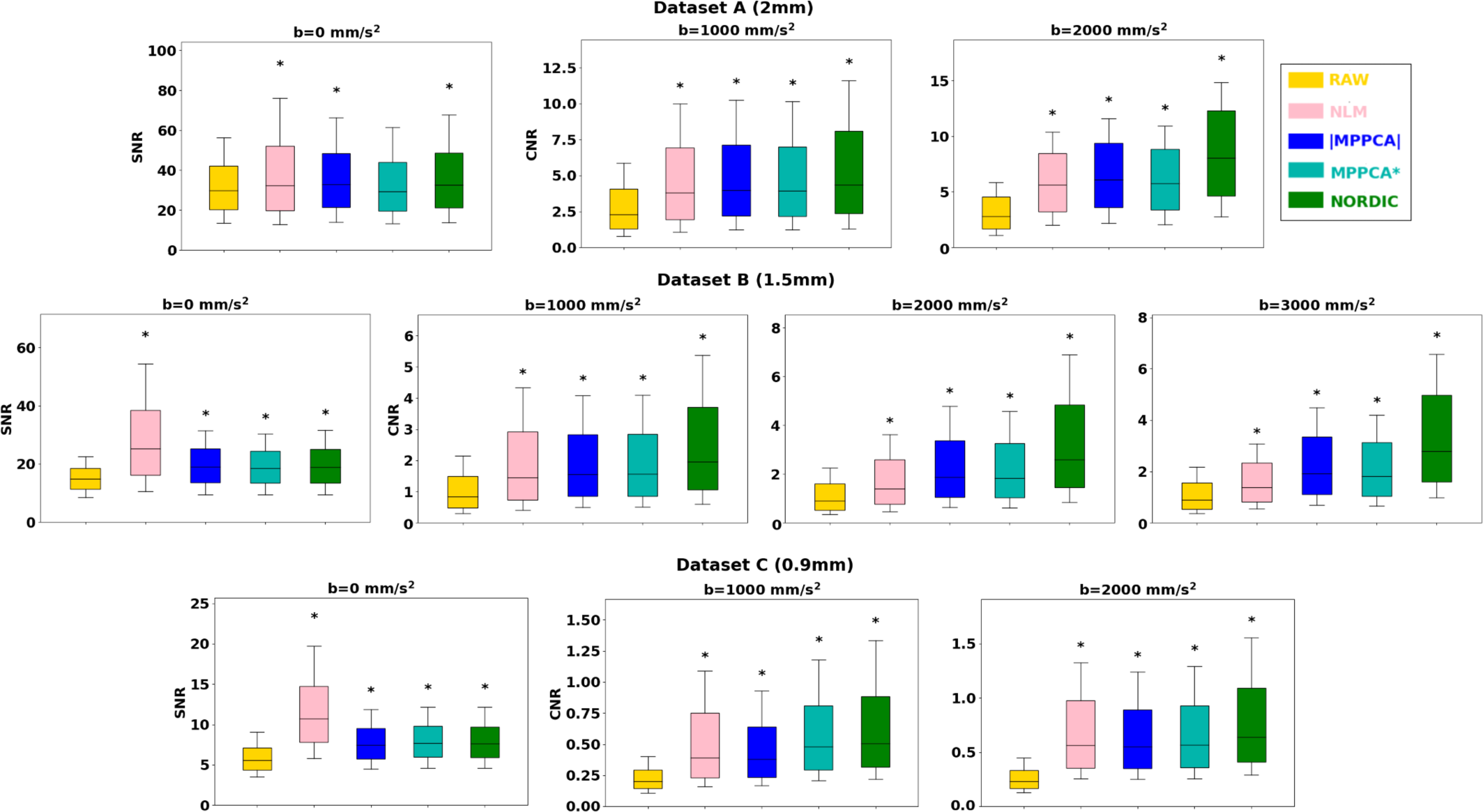
Voxel-wise Signal-to-Noise Ratio (SNR) and Angular Contrast-to-Noise Ratio (CNR) in raw and denoised data. Boxplots correspond to the distribution of these metrics in the whole brain. Top row: Dataset A (2mm), Middle row: Dataset B (1.5mm), Bottom row: Dataset C (0.9mm). Asterisks indicate statistical significance of pairwise Wilcoxon rank-sum tests between SNR/CNR values for raw data vs each denoised approach (Bonferroni corrected for multiple comparisons).

We subsequently looked into the spherical harmonic spectrum of the signal and explored how different denoising methods perform across different angular frequencies. Figure 3 demonstrates changes in the power of spherical harmonic coefficients in the denoised data compared to the raw data. For each harmonic order and each shell, the % change of power of the denoised data in white matter compared to the respective power obtained for raw data is plotted (similarly for grey matter is presented in Suppl. Figure 3). Compared to raw data, all methods removed a high proportion of the power at higher harmonic orders (*L* ≥ 4). At low orders and low SNRs, we can see a different pattern between magnitude-based and complex-based denoising. At *L* = 0, complex denoising reduces more power than magnitude-base denoising (e.g. see Dataset C in Figure 3), while at *L* = 2 they show an increase in power (which is more noticeable as the SNR reduces). This behaviour is also observed when comparing the magnitude- and complex-averages, respectively. This reduction of *L* = 0 power (flat signal) and increase in *L* = 2 power (unimodal anisotropic signal) with complex-domain approaches can be associated with the signal recovered by pushing lower the noise-floor (see Suppl. Figure 4). Indeed, noise floor can rectify the signal in very anisotropic regions and this signal can be recovered by denoising in the complex domain (see also next section). It is important to highlight here how crucial the choice of a gold standard can be in interpreting such results and how different the interpretation can be when using a biased average (magnitude domain).

**Figure 3.**
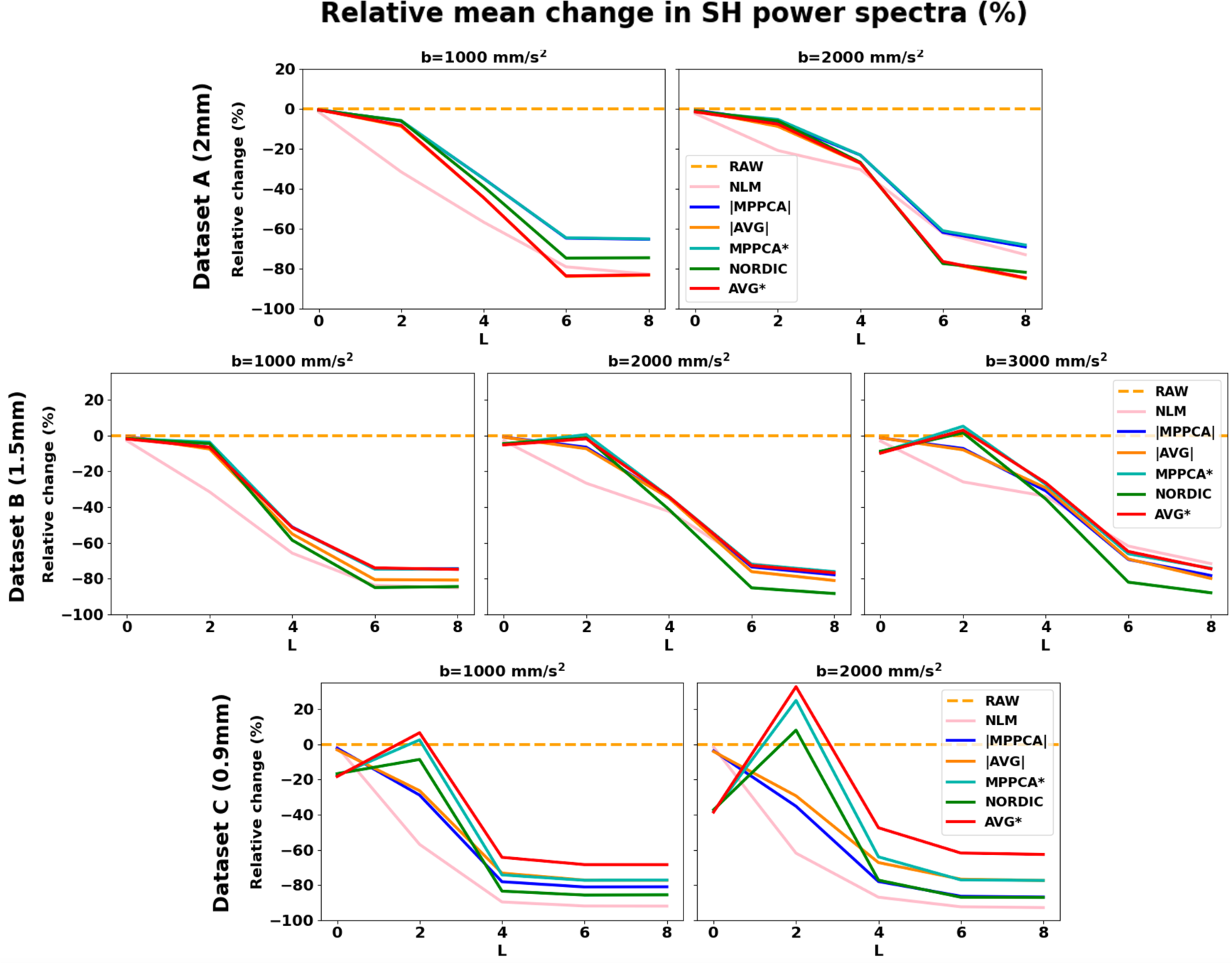
Power spectra of Spherical Harmonics. Mean relative change (%) in the white matter spherical harmonics (SH) power after denoising, with respect to the corresponding power for the raw data. Results for even harmonic orders L = 0 to L = 8 are presented. Top row: Dataset B (1.5mm), Bottom row: Dataset C (0.9mm).

The spatial pattern of the power spectra for different frequencies can be observed in the exemplar brain maps of Suppl. Figure 5 for Dataset C. It can be seen 1) a spatially-homogeneous power reduction at *L* = 0 by complex denoising (and a strong spatial smoothing effect in NLM); 2) an increment of power at *L* = 2 by complex denoising in the white matter; and 3) a considerable reduction of power in the whole brain at *L* = 4 by all denoising methods compared to raw data (a relatively uniform *L* = 4 map prior to denoising, becomes concentrated mostly in white matter areas after denoising).

### 3.2. Dealing with noise floor

We first explored the type of information that is removed during denoising. Figure 4 shows difference maps estimated between the raw magnitude data and the denoised images within the brain, and their corresponding histograms, for Datasets B and C. Ideally, these differences should arise only from thermal noise, which is spatially randomly distributed in nature, leading to a zero-mean Gaussian distribution of the intensity values. This is the behaviour that, a priori, can be observed for magnitude MPPCA and that has been extensively reported in previous studies as a sign of good denoising (e.g., (Veraart et al., 2016b)). However, zero-mean Gaussian noise properties only hold in the complex domain. In the magnitude domain, a Rician/non-Chi-squared signal distribution is instead expected, which induces a noise floor. Hence, denoising that induces a zero-mean (or negative mean, as in NLM) difference map with respect to the magnitude raw data is indicative of retaining (or even increasing, as in NLM) the noise floor. We can observe that denoising in the complex domain leads to a non-zero (positive) mean-centred distribution of differences (see histograms of Figure 4)), meaning that denoised signal intensities are on average lower than in raw data. This is commensurate with the hypothesis that the signal rectification caused by the noise floor is reversed and the associated signal elevation is removed.

**Figure 4.**
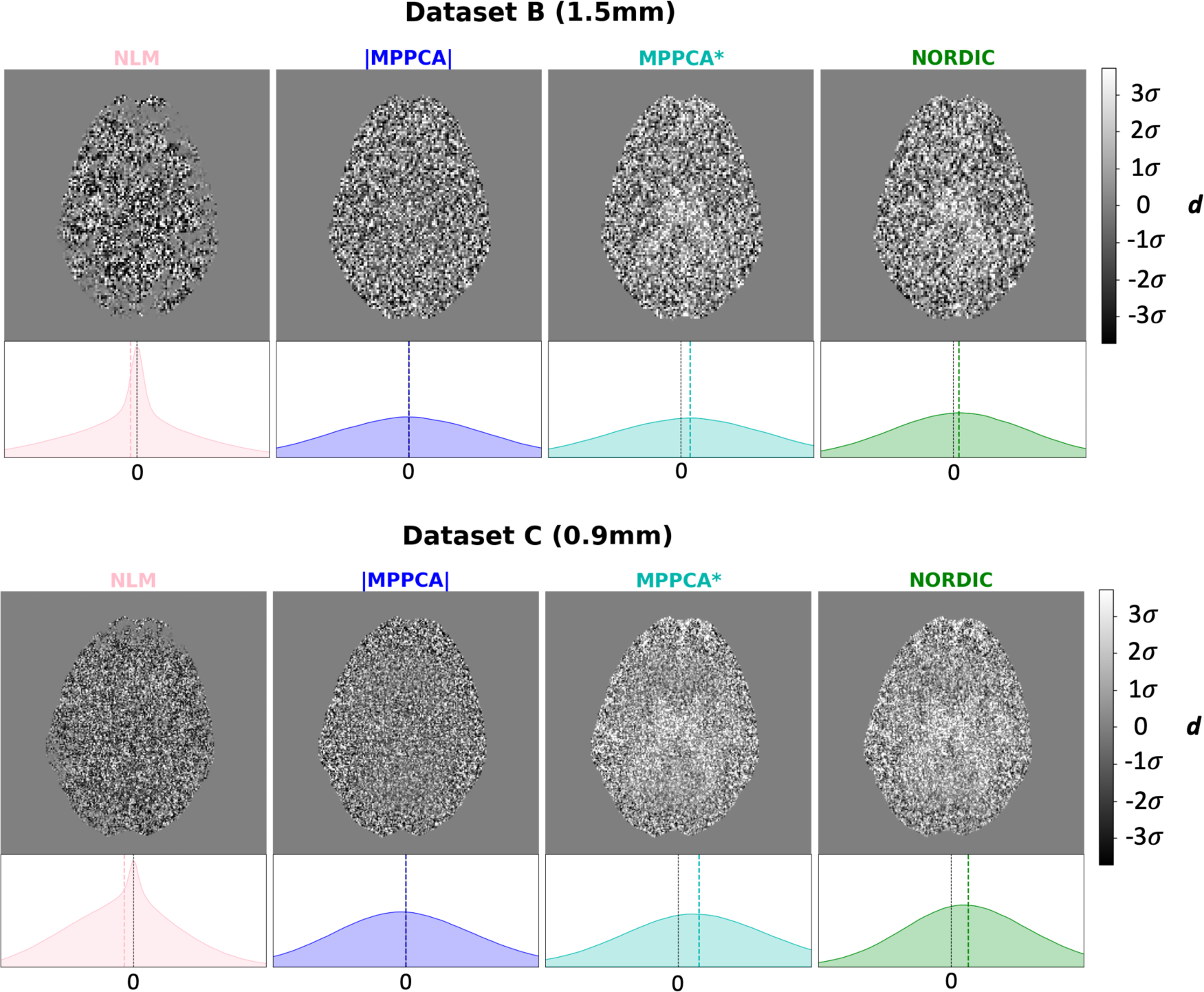
Difference between raw magnitude and denoised data - The difference Δ, where Δ = S_RAW_ −S_denoised_ for each voxel, is represented for each method in an exemplar axial slice, and its corresponding histogram of values within the brain is shown. In the histogram, the vertical black dashed line indicates Δ = 0, while the coloured dashed line indicates the mode of the actual distribution of the differences. For each dataset, all plots are displayed in the same range of values (as a function of the standard deviation σ of the distribution, indicated by the colorbar). The orange arrow points to the ventricles, where the effect of the noise floor is more noticeable and the difference between non-denoised and complex-denoised data becomes higher (i.e., the signal in the ventricles of the non-denoised is higher as it gets rectified).

To explore this further, we looked at the signal at places where noise floor rectification is anticipated, as shown in Figure 5. On the left, the signal intensity distribution from voxels in the ventricles across all volumes is shown for each method in Dataset C (0.9mm). The signal is maximally attenuated in these CSF-filled regions, so what is effectively depicted is a proxy to the distribution of noise, before and after denoising. The histograms of all denoised datasets show an evident reduction of the signal variance (more restricted range of values), as expected and in agreement with the SNR/CNR trends shown before. The mode of this distribution is roughly equivalent to the raw magnitude data for all denoising methods that operate in the magnitude domain. However, the mean of the noise distribution is reduced for MPPCA*, NORDIC, and AVG*, consistent with that denoising in the complex domain reduces the noise floor.

**Figure 5.**
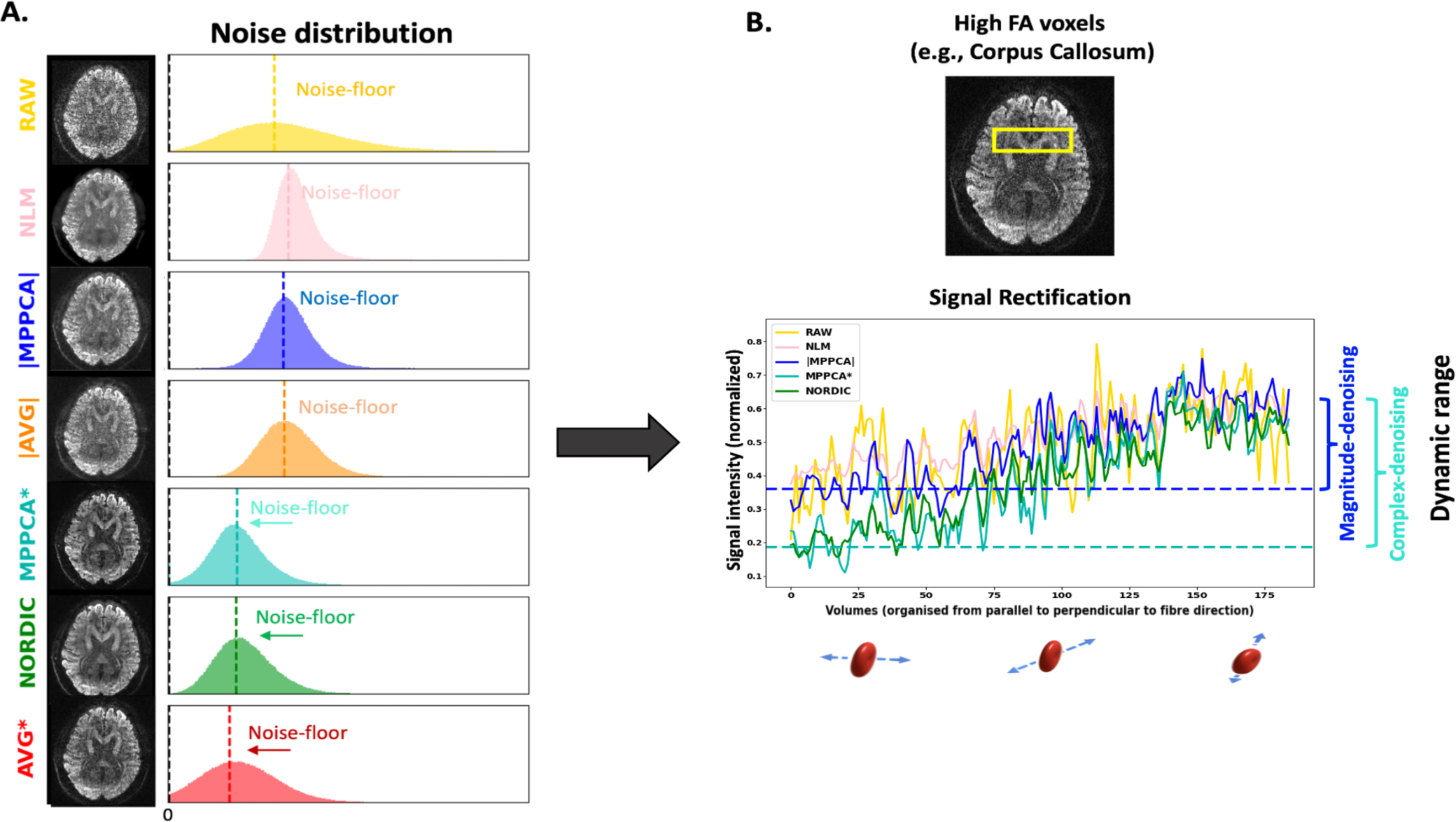
Noise-floor effects and signal rectification (Dataset C) – A) Histogram of the signal intensity values in the ventricles obtained from the different denoised datasets. B) Signal intensity values in an example high FA voxel (FA>0.8) from the Corpus Callosum. Signal for different diffusion-sensitising volumes has been ordered so that gradient orientations are from parallel to perpendicular to the primary fibre orientation (DTI v_1_) of AVG*, i.e. approximately lowest to highest signal intensity. To aid visualisation, dashed lines in panel B represent the median value of the first points of representative denoised signals (*complex and magnitude MPPCA*). These reflect the lowest signal (i.e. parallel to the main fibre orientation) for that WM voxel in each case. An indication of the corresponding dynamic range is also shown as the difference between these lowest signals against the maximum signal in each case (median of last 15 points for each case, reflecting signal roughly perpendicular to the main fibre orientation).

On the right plot of Figure 5 we show the signal of an example anisotropic voxel in the mid-body of the Corpus Callosum in Dataset C (more examples shown in Suppl. Figure 6). The raw and different denoised dMRI signals for that voxel have been sorted based on the alignment of the corresponding diffusion-sensitising gradient direction and the primary fibre orientation in this voxel (given by the DTI primary eigenvector of AVG*). Hence, the signals from directions parallel to the primary fibre orientation are presented first, and signals perpendicular to it are presented last. Consequently, the first signals are maximally attenuated, while the last signals are minimally attenuated. Before denoising (yellow line), the maximally attenuated signals have been rectified by the noise-floor and appear roughly similar to the minimally attenuated signals (i.e., small dynamic range). This behaviour is preserved when denoising in the magnitude domain. On the other hand, denoising in the complex domain increases the dynamic range of the signal and mitigates the effects of the noise floor.

### 3.3. Effects of patch-based denoising on spatial resolution

We subsequently explored the performance of the different denoising methods (all patch-based) with regards to spatial resolution. Figure 6 shows the estimated resolution in the images before and after denoising compared to the nominal resolution (dashed line). DTI model residuals have been used to estimate smoothness in the data (version with DKI model residuals is presented in the Suppl. Figure 7, showing similar trends). As noticed in the raw data, due to the point-spread-function blurring, resolution along the phase-encoding direction (y-axis) is always smaller than nominal, while in the frequency encoding direction (x-axis) it is very close to nominal, as expected. We can observe that all denoising methods incur a resolution penalty.

**Figure 6.**
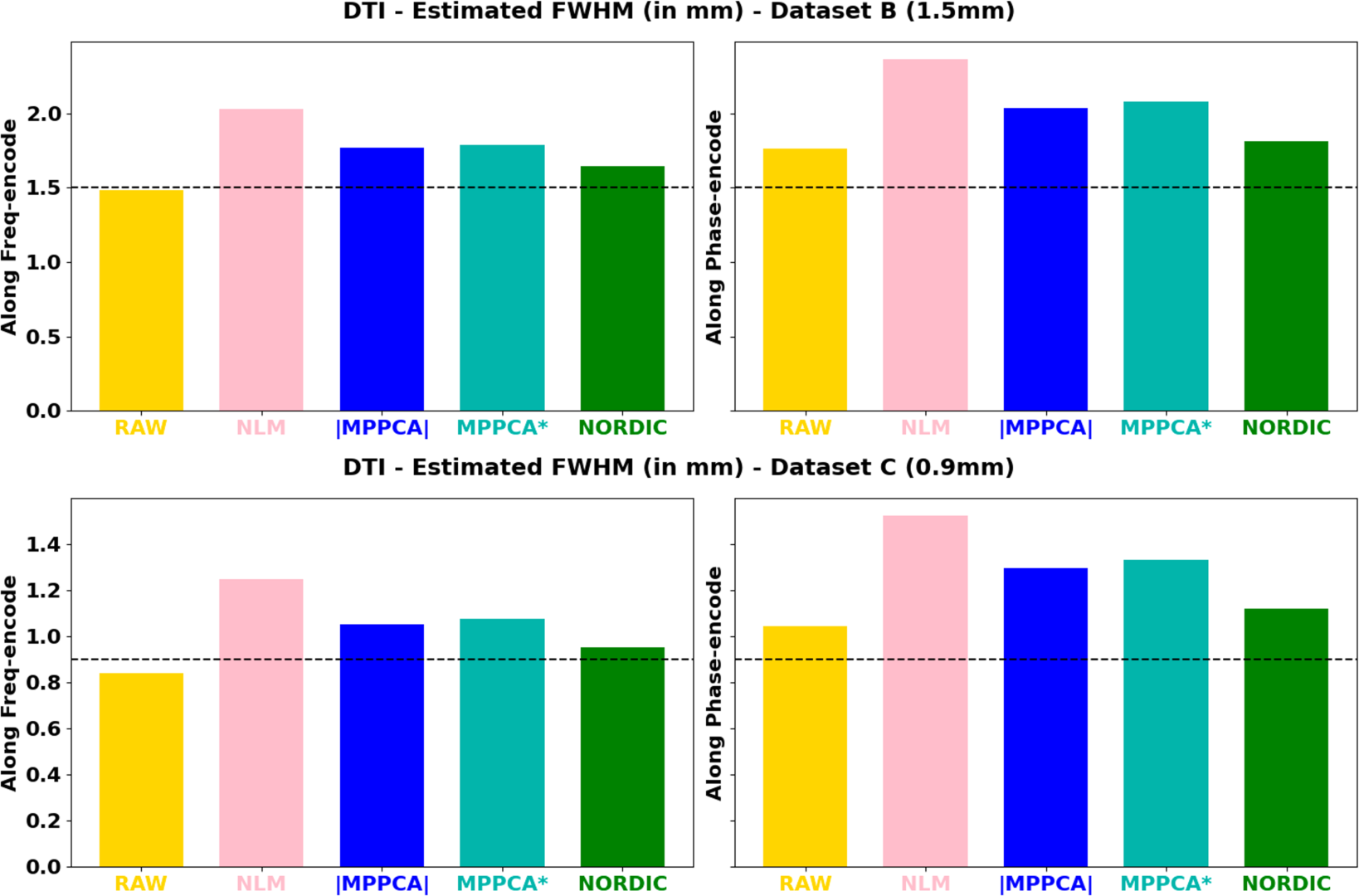
Estimated voxel resolution for the denoised and non-denoised datasets, along the frequency (acquisitions along x-axis) and encode (acquisitions along y-axis) directions - Top row: Dataset B (1.5mm); Bottom row: Dataset C (0.9mm). Dashed line: acquisition nominal resolution.

Table 1 quantifies that further and shows the % resolution change along the frequency and phase encoding axis of each method with respect to the raw data. The resolution penalty ranges from 5 to 45% and is worst for NLM that causes more substantial blurring, agreeing with other studies (e.g., (Mishro et al., 2021)). NORDIC keeps closer to the original resolution overall (penalty in the order of 5 to 15 %), followed by MPPCA (between 20 to 30 %).

**Table 1.**
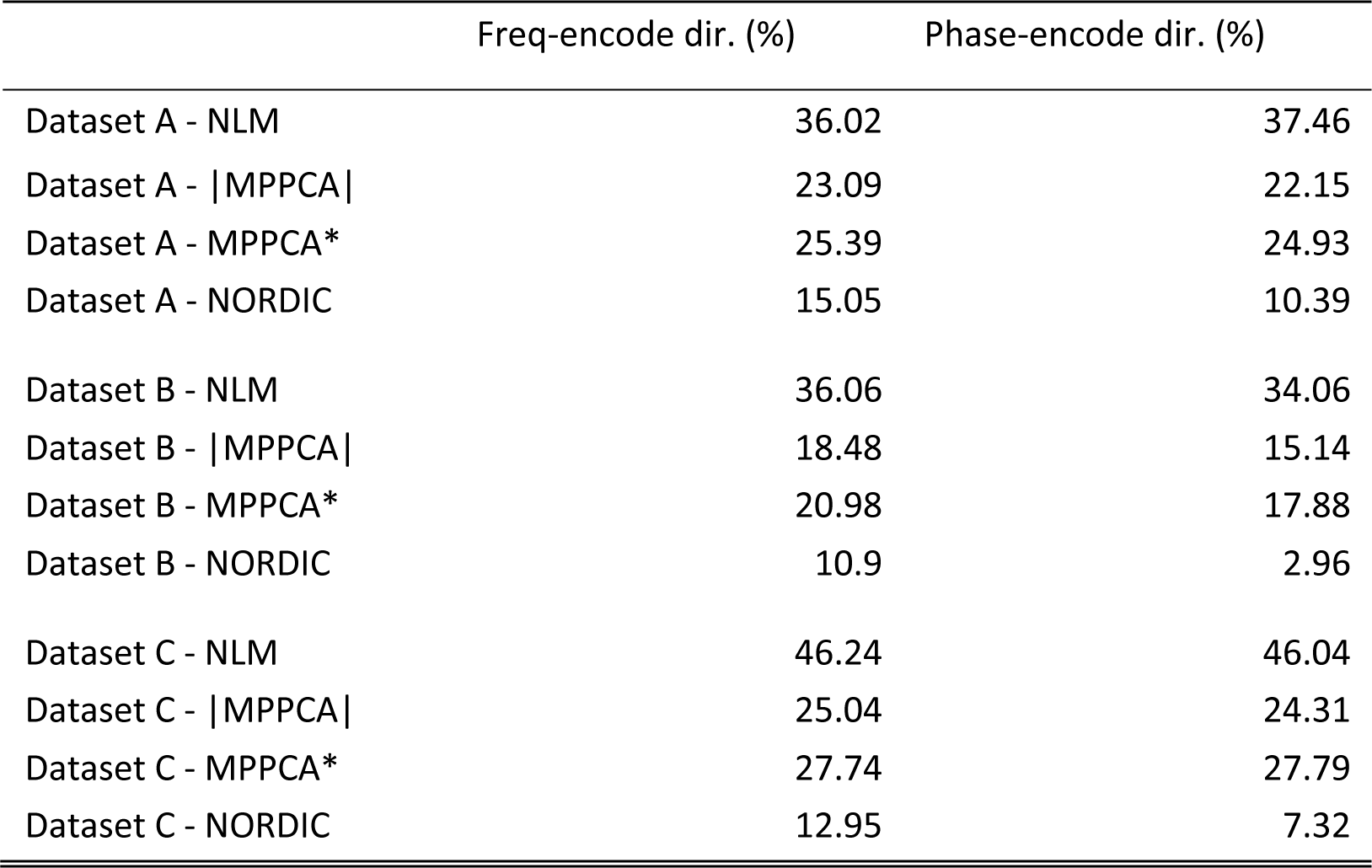
Spatial resolution loss induced by each method in each dataset, quantified as the percentage difference between the denoised and raw data estimated resel (FHWM), i.e. 100 · (FWHM_denoised_ − FWHM_RAW_)/(FHWM_RAW_)

These results were obtained using the default implementation of each denoising algorithm, therefore reflecting a different default patch-size. Both MPPCA and NORDIC implementations adjust the patch size by the number of volumes *N*. In MPPCA *dwidenoise* implementation, an isotropic patch *P* is automatically adjusted, such that *P* is the smallest odd value that exceeds the number of DW images *N*, i.e. *P*^3^ ≥ *N*; in NORDIC, *P* is adjusted to maintain an 11:1 ratio between voxels in the patch and *N*, i.e. *P*^3^ ≥ (11 ∗ *N*). Therefore, to better understand the previous results, we explored performance across a range of patch-sizes for the different approaches. Figure 7 shows the FWHM estimated at different patch-sizes for the different methods. Asterisks show the default patch size in each implementation for each dataset. We also performed CNR calculations for each patch-size to accompany potential spatial resolution penalties with median gains in signal quality of the white matter (Figure 8). Taken together these two figures suggest that: 1) regardless of the nominal resolution, larger patch sizes induce smaller penalties in spatial resolution after denoising, converging to the non-denoised data if the patch is big enough (this patch size will depend on the dataset and the algorithm), and 2) there is an inverted U-shape in the CNR gains suggesting that increasing the patch size after a certain point reduces denoising CNR gains. The optimal patch-size varies and the higher the b-shell (or the noisier the data is), the higher the patch size needed to provide maximum gains of CNR. 3) The patch size used in the default NORDIC implementation seems to be closer to optimal in terms of minimising spatial resolution penalty and maximising CNR gain, compared to the default MPPCA implementation used here. Notice that NLM was not included in these plots, its performance was apparently suboptimal for patch sizes larger than 3^3^ (the default), as high levels of blurring were introduced (see Suppl. Figure 8).

**Figure 7.**
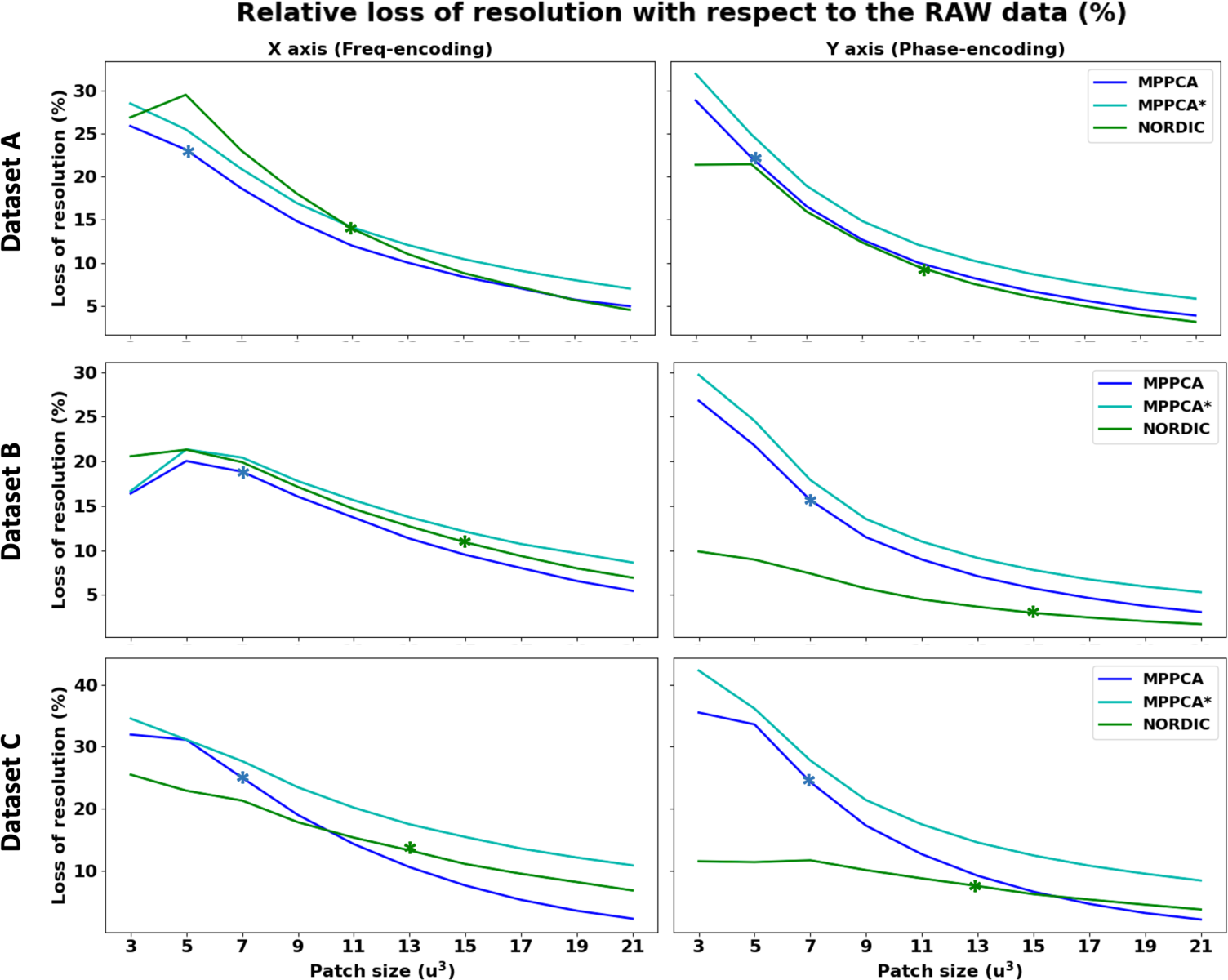
*Percentage* loss of resolution (given by the estimated FWHM) as a function of the patch-size used in the denoising algorithm. Asterisks indicate the default patch-size of each algorithm. Top: Dataset A (2mm); Middle: Dataset B (1.5mm); Bottom: Dataset C (0.9mm).

**Figure 8.**
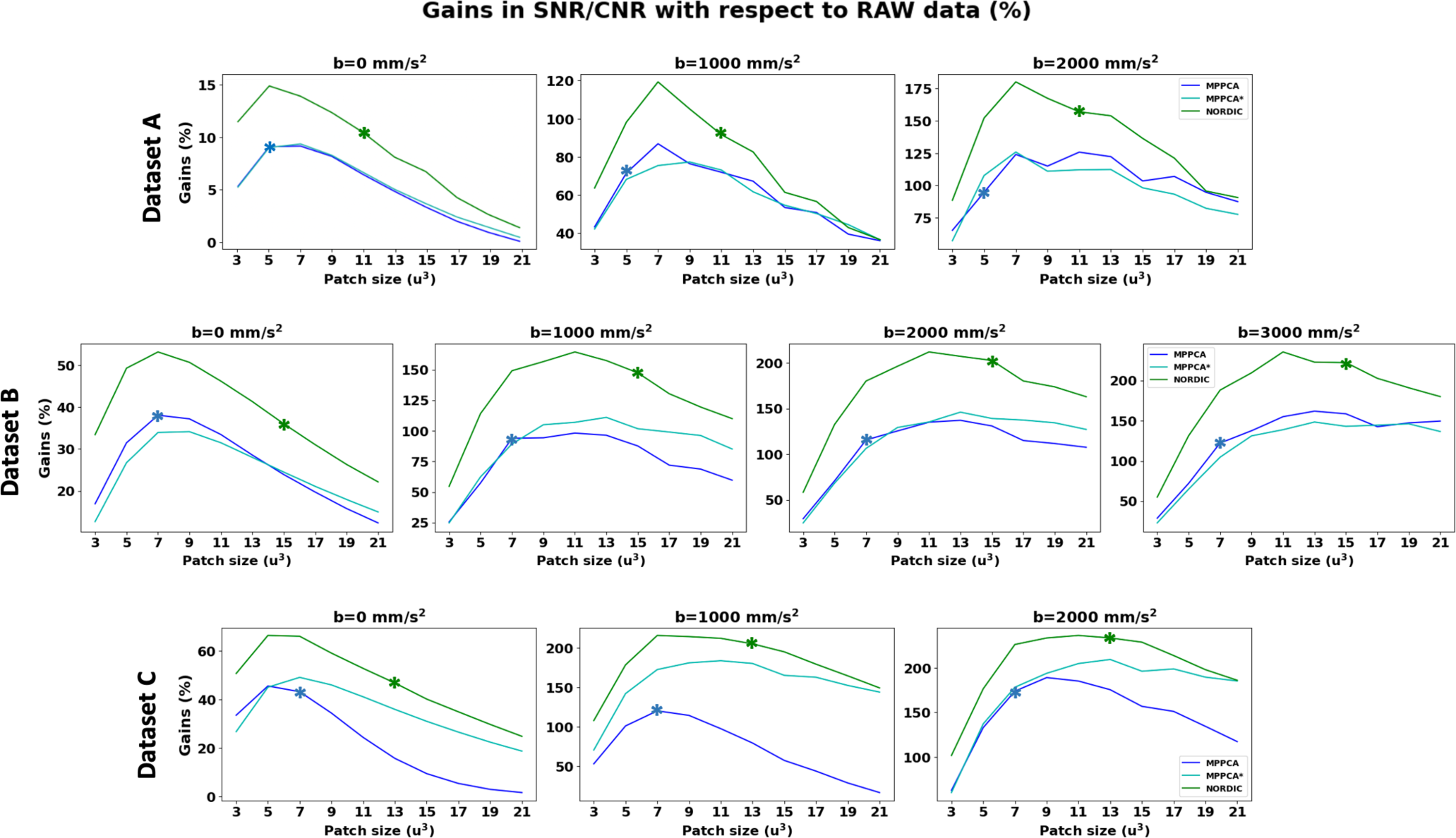
*Median* gains in signal quality in the white matter at different b-shells with respect to the non-denoised data (per columns) as a function of the patch-size. Top: Dataset A (2mm); Middle: Dataset B (1.5mm); Bottom: Dataset C (0.9mm). The patch that provides the highest gains in signal quality depends on the resolution and the b-shell.

### 3.4. Gains in precision and accuracy for model estimation

We evaluated whether denoising gains in SNR/CNR (Figure 2) and behaviour against noise floor effects (Figures 3, 5) were translated into downstream improvements in precision and accuracy of model estimates respectively. For the accuracy evaluation, Figure 9A shows the correlation between DTI estimates (FA and MD) obtained from different repeats of denoised and non-denoised data against those obtained using the complex-average (AVG*), assuming the latter represent a gold standard. Both MPPCA* and NORDIC show substantial improvements across all tested datasets with respect to the non-denoised data. Of interest is the behaviour of |MPPCA| that, particularly for MD in a very noisy scenario (Dataset C), does not converge to the gold standard and offers reduced benefits compared to non-denoised data. This is indicative of the noise floor effects, discussed in the previous sections, which are reduced when denoising in the complex domain, but remain high when denoising in the magnitude domain.

**Figure 9.**
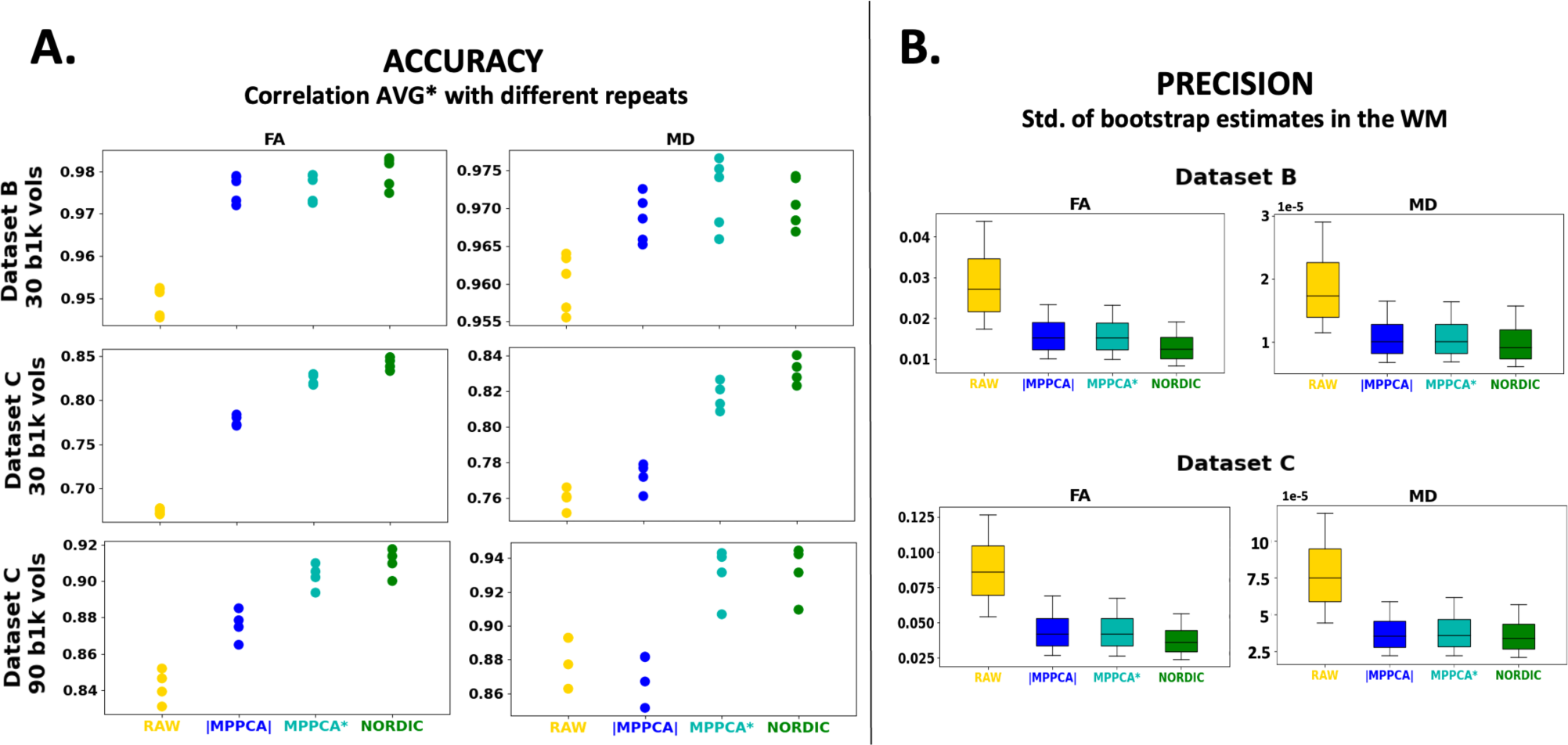
Precision and accuracy in DTI model estimates. A) Accuracy in FA and MD from non-denoised and denoised data is evaluated against FA and MD estimated using the complex average (AVG*) of multiple repeats. Plots show correlation values within WM. Different dots represent one of the repeats (5 for dataset B and 4 for dataset C). This is shown for different exemplar scenarios: Dataset B (1.5mm resolution) with 30 b=1000 s/mm^2^ volumes (top); Dataset C (0.9mm resolution) with 30 b=1000 s/mm^2^ volumes (middle); Dataset C with 90 b=1000 s/mm^2^ volumes. B) Precision in FA and MD evaluated using wild residual bootstrap. The uncertainty (inverse precision) in the FA and MD estimates is calculated as the standard deviation across 250 bootstrap samples.

For the precision evaluation, Figure 9B represents the uncertainty of DTI estimates across two different datasets, calculated using bootstrapping. The trends demonstrate that the denoising benefits in SNR/CNR shown before (Figure 2) translate to uncertainty decrease / precision increase in both FA and MD, compared to non-denoised data. Improvements are at least two-fold in most cases following the almost two-fold average CNR increase of the b=1000 data in Datasets B and C after denoising.

### 3.5. Convergence at the high-SNR limit

We assessed convergence at the high-SNR limit by evaluating benefits across different SNR regimes. As we showed earlier, our low-resolution dataset (A) has the highest SNR/CNR, and our ultra-high-resolution dataset (C) has the lowest SNR. We, therefore, explored how denoising gains vary with our three datasets used to probe performance differences at different SNR regimes. In Figure 10 (top) we show the gains in the median SNR and angular CNR by denoising compared to raw data, as we move from low SNR (dataset C) to medium SNR (dataset B) and high SNR (dataset A). NORDIC provides the largest benefits overall, nevertheless, all methods show greater benefits for the lowest SNR dataset and for the highest b-value; and the smallest benefits for the highest SNR dataset and lowest b-value. The bottom figure shows a similar pattern of tractography performance convergence evaluated against tractography results obtained using the complex average data (AVG*). Agreement with the gold standard improves substantially after denoising compared to raw data in the low SNR case. Denoising has effectively no effect in higher SNR data (in fact NLM slightly decreases performance compared to raw data). Taken together, these results suggest convergence of the denoising performance at the high-SNR regime, with complex denoising approaches performing better overall for noisier data.

**Figure 10.**
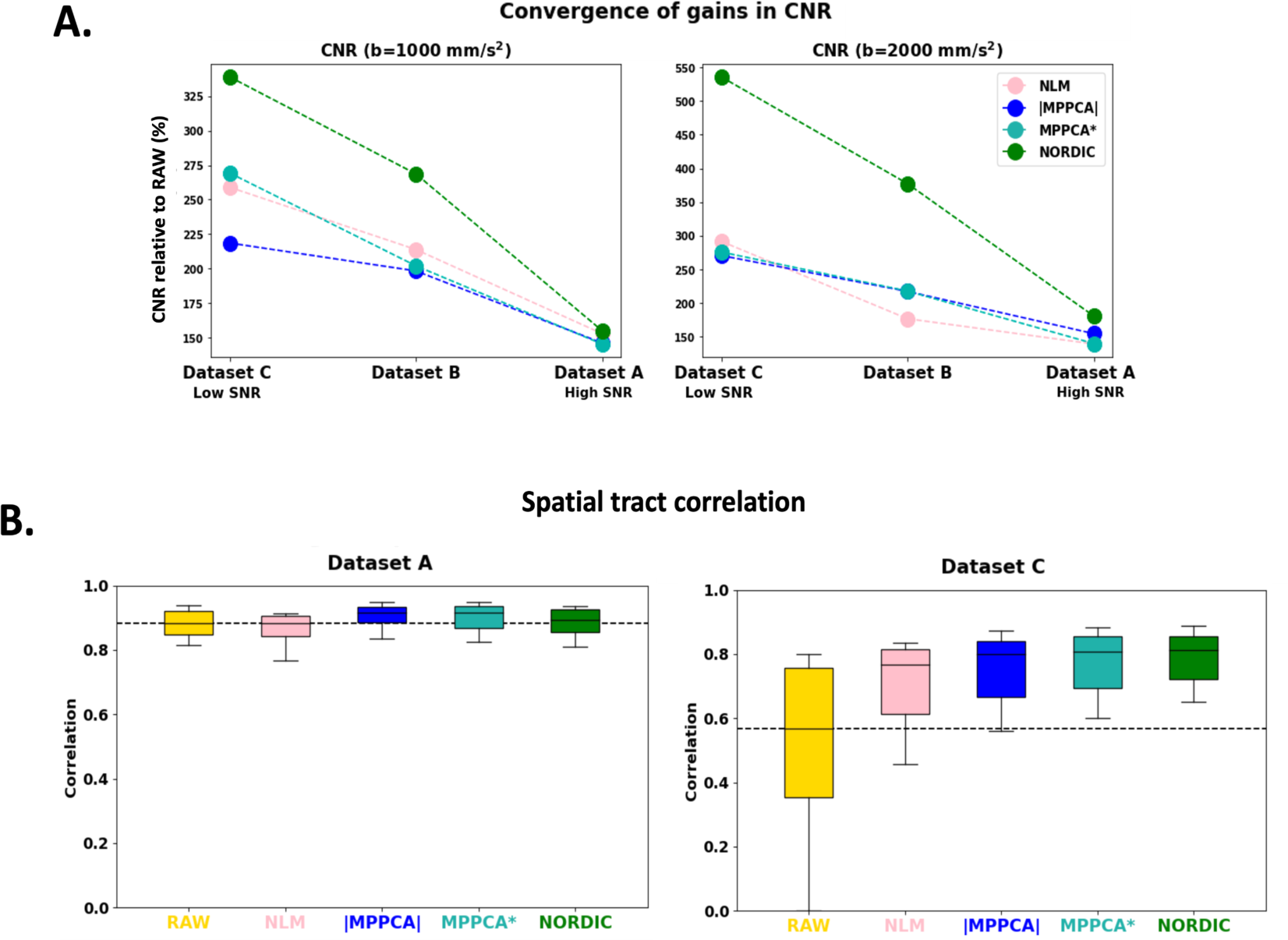
Convergence of performance with higher SNR. Gains in denoising performance for different SNR examples (low, medium and high SNR) are presented. A) Percentage change of median CNR (per b-shell) after denoising with respect to the angular CNR of the raw data, and B) Spatial tract correlation across 42 bundles with the tracts obtained from the complex average for the low resolution/high SNR data (left) and the high resolution/low SNR data (right).

### 3.6. Denoising gains for SNR-limited applications

Improvements offered by denoising can open opportunities that only bespoke setups can provide, such as allowing usable data at ultra-high spatial resolution; or reducing scan time, as the increased SNR per volume reduces the need for sampling many volumes or multiple repeats. In this section, we explore the feasibility of these two applications.

#### 3.6.1. Benefits for pushing spatial resolution

In the previous figure, we showed that denoising had a large effect on tractography performance for the ultra-high resolution dataset. Acknowledging that a 0.9mm isotropic dMRI dataset acquired with a conventional PGSE sequence in a clinical scanner is borderline unusable in most cases, we considered tractography using such data as a challenging scenario. We, therefore, explored what difference denoising methods can make in a demanding application.

We first looked at sensitivity in estimating fibre orientations. Figure 11 demonstrates how multi-way crossings in the Centrum Semiovale (a region where most of the voxels are expected to have at least two fibre bundles crossing) can be recovered after denoising in the 0.9mm data. NORDIC provides overall higher rates of detection of secondary and third fibres at 0.9mm resolution, although both MPPCA algorithms also show considerable improvements over raw data. As previously shown (Figure 9), precision is also improved by denoising (here uncertainty in fibre orientation is assessed by MCMC on the ball & sticks model, focusing on a region where most voxels are expected to exhibit a single fiber, the midbody of the Corpus Callosum).

**Figure 11.**
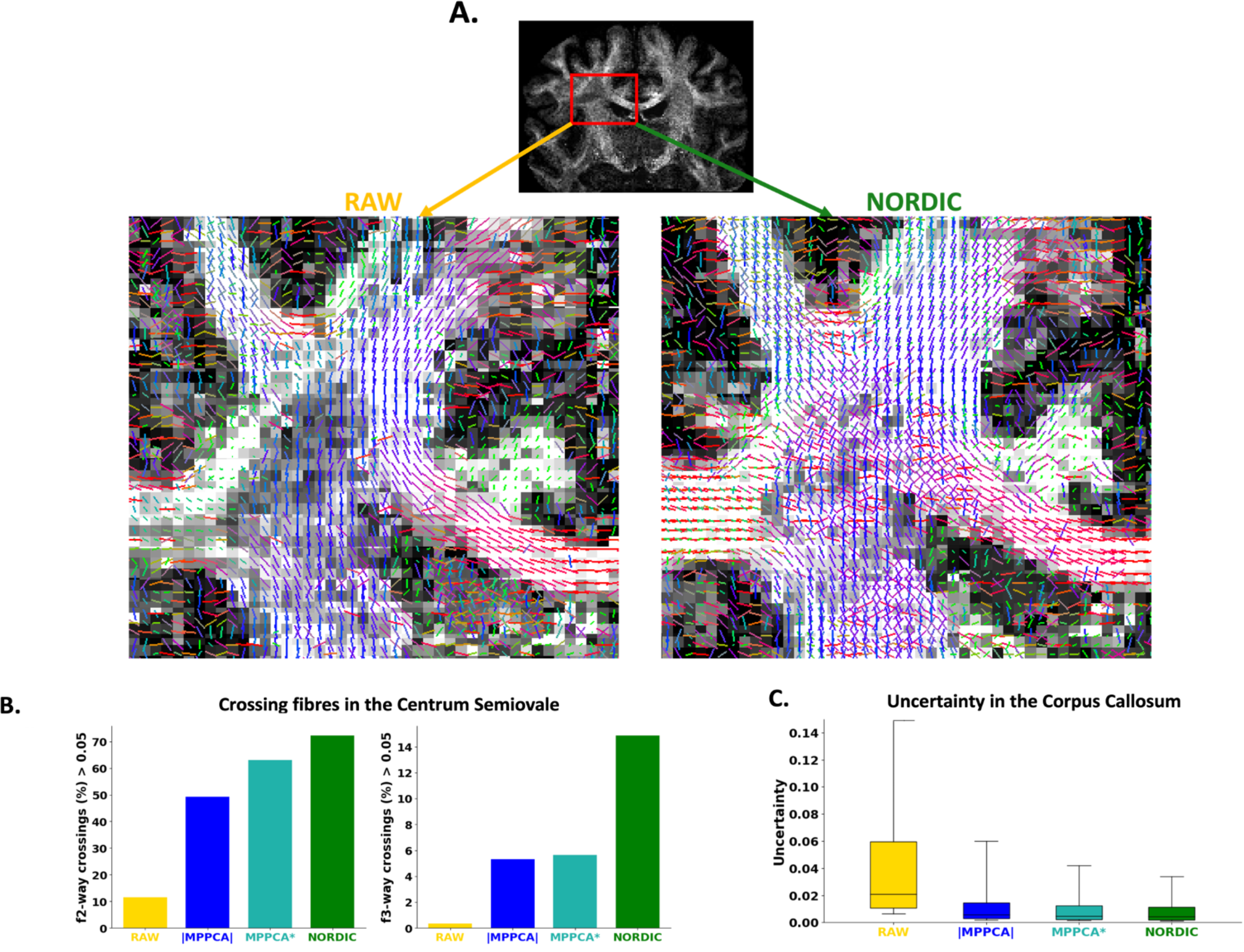
A comparison of modeling performance between RAW and denoised data (Dataset C - 0.9mm) - Top: Example of fibre crossings detected in the Centrum Semiovale in the RAW and NORDIC-denoised data. Bottom left: Rate of crossing detection in the Centrum Semiovale. Bottom right: First fibre uncertainty measured in the voxels of the Corpus Callosum.

We subsequently looked in greater detail at tractography success at this low SNR regime. Figure 12 demonstrates the tractography of 42 major white matter bundles (Warrington et al., 2020) in the raw vs denoised data. A number of tracts could not be reconstructed at all using the raw data (seven bundle tracts in total, including the left Acoustic Radiation, left and right Corticospinal Tracts, right Fornix, Middle Cerebellar Peduncle, and left and right Superior Longitudinal Fasciculus 1). Denoising allowed good reconstruction of all considered WM tracts, apart from the Middle Cerebellar Peduncle in MPPCA and NLM, and of all 42 tracts for NORDIC. As shown in Figure 10B, all denoising methods improved the agreement of tractography results with the HCP Atlas, compared to the raw data. Denoising in the complex domain also showed the highest improvement and was closest to the tracts reconstructed with the higher-SNR, 2mm dataset.

**Figure 12.**
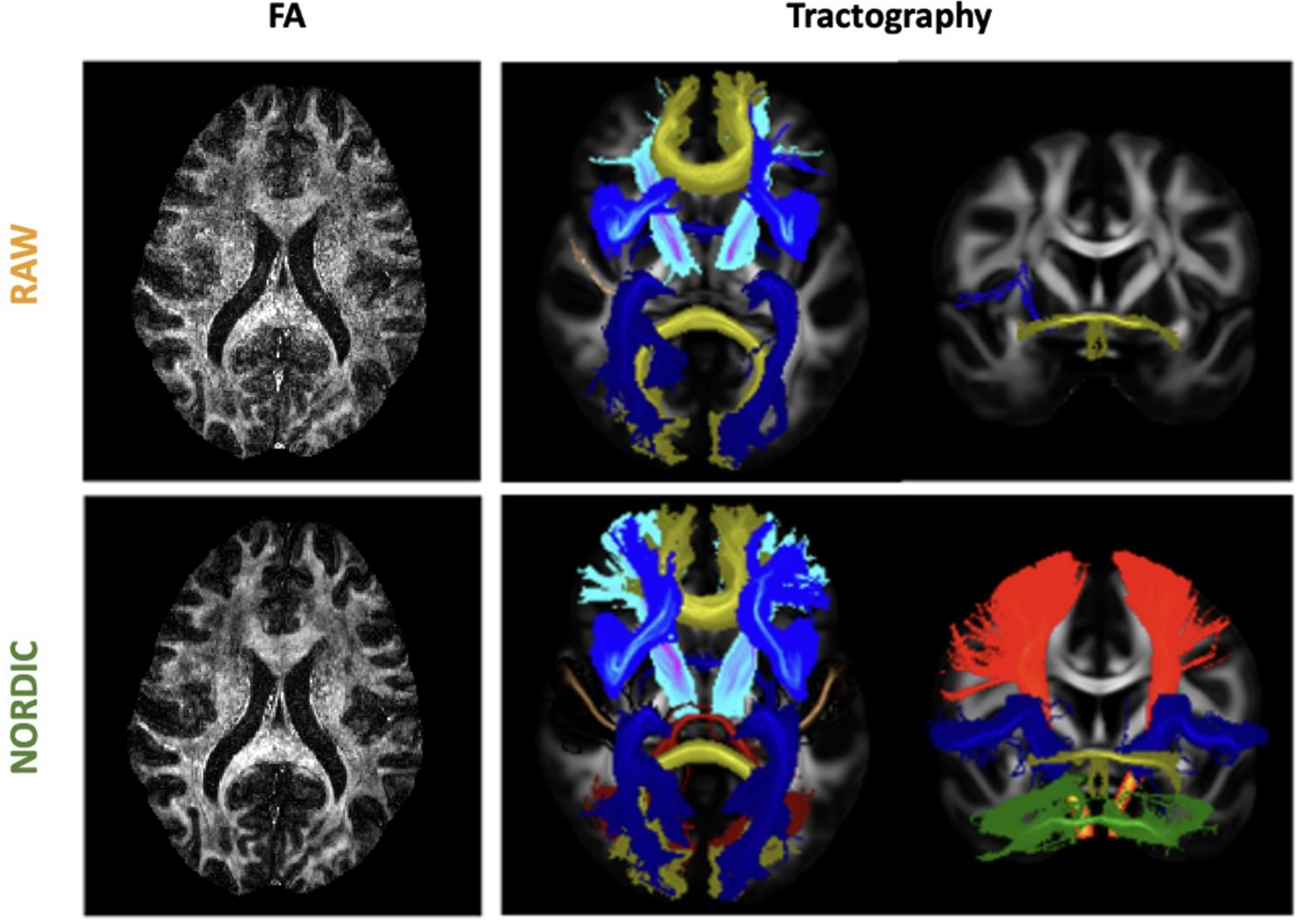
Examples of FA maps (left) and Maximum Intensity Projection (MIP) of tracts reconstructed (right) by RAW and NORDIC-denoised data at 0.9mm (Dataset C). At this resolution, denoising allows to recover multiple bundles that were missed in the non-denoised version.

#### 3.6.2. Benefits for reducing scan time

Denoising inherently aims to improve the effective SNR per sample. This opens the question of whether scan time can be reduced, i.e., whether acquiring fewer denoised volumes is equivalent to acquiring many noisy volumes. Figure 13 shows a qualitative comparison for fractional anisotropy maps, before and after NORDIC denoising from Dataset C and how these compare against maps from multiple averages Figure 13 (top) and from considerably longer scan times in a single-repeat acquisition Figure 13 (bottom). These illustrate how denoised, subsampled data (either fewer directions or fewer repeats) improved agreement to the full dataset (more directions or repeats, respectively) compared to maps obtained from similarly subsampled but non-denoised data.

**Figure 13.**
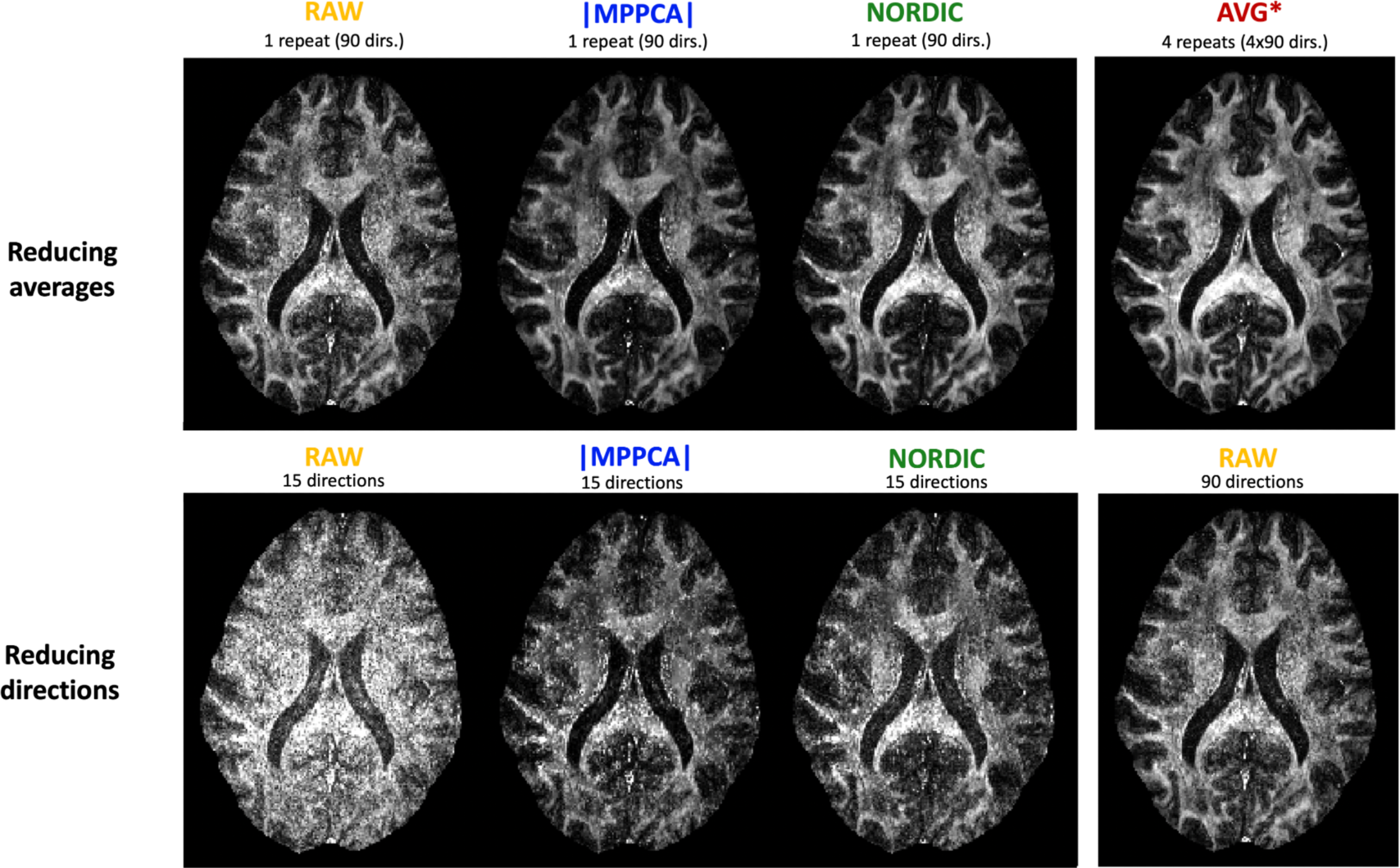
Qualitative FA maps comparisons (Dataset C - 0.9mm). Top row: Denoised single-repeat data vs. complex multiple-averages. Bottom row: A low number of volumes from denoised data vs high number of volumes from RAW.

To quantify these improvements, we calculated the sum of squared differences and spatial correlation between FA and MD scalar maps in WM obtained from subsets of the original data against the FA and MD maps obtained from the complex average of multiple repeats of the original data. Figure 14 shows the agreement between FA and MD maps in Dataset C (AVG* used as a reference), where MPPCA* and NORDIC converge faster to the reference than the non-denoised data. We can observe that with complex denoising even with ∼ 30 − 35% of the number of volumes we can achieve similar or better agreement to the gold standard average as the full non-denoised dataset. These results also suggest that denoising with one of the complex approach as low as 4/6 of the considered datapoints, one can achieve correlations of ∼ 0.9 against the gold standard; this is equivalent to about 14 minutes of 3T scanning for a 0.9mm isotropic multishell dataset.

**Figure 14.**
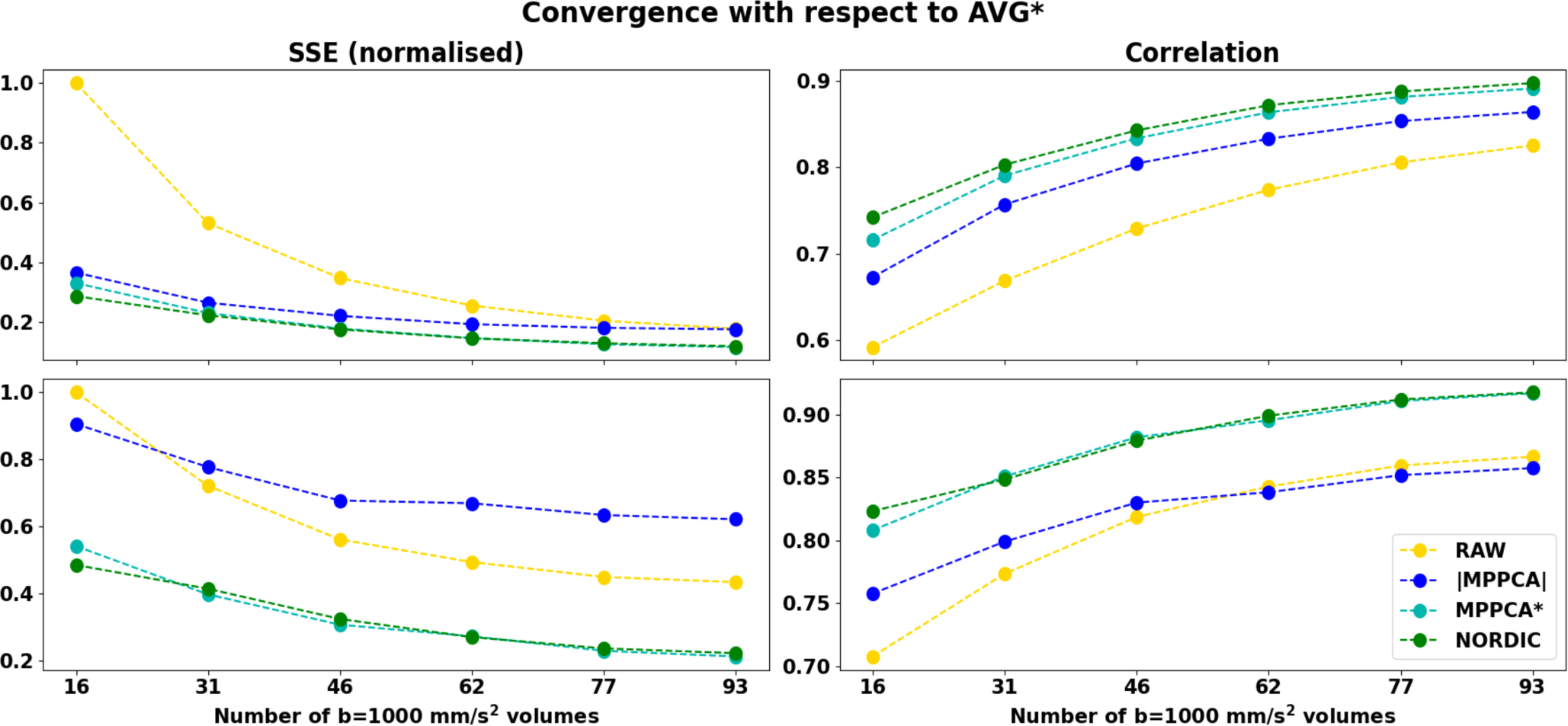
Convergence to the complex multiple-averages (reference) assessed by the Sum of Squared errors (SSE) (left column) and Pearson correlations (right column) in DTI model estimates of subsets from Dataset C (0.9mm). The x-axis represents increasing fractions of volumes used to fit the DTI tensor, corresponding to 1/6, 2/6, …6/6 of the whole dataset that contains 93 b=1000 s/mm^2^ volumes. Top: Fractional Anisotropy correlations. Bottom: Mean Diffusivity.

We also noticed that approaches that perform denoising in the magnitude domain seem to converge to a different reference compared to complex denoising approaches. We performed the same comparisons using the magnitude average as a reference (Suppl. Figure 9) and demonstrated how magnitude-based methods converge to this reference rather than the complex average. The deviations are more evident when looking at the MD, which is expected to be more affected by the noise floor effects, as described in the previous sections. To confirm this, we assessed agreement in WM regions that are expected to have lower noise floor effects (regions that have low FA, i.e. lower attenuation). Doing so for FA*<*0.2-0.3, we observed a better agreement of complex denoising methods against the magnitude average (results not shown). Taken together these results demonstrate the feasibility of reducing scan time using denoising approaches and the importance of selecting the correct reference to compare performance against in an unbiased manner.

## 4 DISCUSSION

We have revisited the evaluation of diffusion MRI denoising approaches. We acquired a novel dataset (both magnitude and phase) that spans a range of SNRs and resolutions and that we publicly release in OpenNeuro. We used this dataset to assess the performance of a number of patch-based denoising methods both in the magnitude and complex domains. We reconsidered what a denoising algorithm should do and proposed a number of criteria, ranging from gains in raw signal quality to enabling SNR-starved applications. We also provide data and code to obtain a gold standard based on complex averaging of multiple repeats against which we compared denoising approaches; we showed that there is a range of quantitative agreement against this gold standard. We also highlighted differences in interpretation and challenges when using a magnitude-average as a gold standard instead of a complex-average. We showed that all denoising implementations reduce noise-related variance as expected, but not always biases from the elevated noise floor. We found that patch-based denoising induces spatial resolution penalties, but its extent can vary depending on the method and the implementation.

Taken together, our results suggest that dMRI denoising in the complex domain is advantageous compared to denoising in the magnitude domain with respect to all of the considered criteria. We found more noise-floor suppression, higher SNR/CNR without strong penalties on spatial resolution, better modelling performance and higher agreement with the gold standard average of multiple repeats when denoising complex rather than magnitude data. This is not surprising; some of the assumptions on thermal noise made by algorithms (e.g., zero-mean Gaussian distribution for the MP law) are better fulfilled in the complex domain. Hence, algorithms that operate in the complex domain (like NORDIC and MPPCA*) could better address noise-floor biases, which are maintained when denoising magnitude data.

We specifically demonstrated that denoising in the complex domain results into a non-zero mean difference map between raw magnitude and denoised data (Fig. 4), with that mean reflecting the noise floor. In the case of an elevated noise floor, denoising that induces a zero-mean difference map is indicative of preserving the noise floor. We further showed that in the spherical harmonic power spectrum reducing the noise floor is reflected as a reduction in power for the DC component, in favour for the power of the L=2 component (Fig. 3); signal which has been rectified by noise floor signal (i.e. relatively constant) gets reduced and replaced by L=2, i.e. anisotropic signal, which agrees with the behaviour demonstrated in Fig. 5B for very anisotropic signal in WM. Note, however, that no approach fully eliminated noise floor (Fig.5A). Nevertheless, this is an important outcome as noise inherently affects both accuracy (bias) and precision (variance); particularly given that reduced precision can be mitigated by acquiring more data, but noise-floor and bias effects cannot be necessarily mitigated by longer scan times. It is not uncommon practice that noise floor effects are either ignored or left to be dealt with at post-"denoising" stage (Koay and Basser, 2006; Koay et al., 2009; Ma et al., 2020a), which is arguably counter-intuitive and suboptimal, as rectified dMRI signal and lost dynamic range cannot be recovered post-denoising. We showed here that denoising in the complex domain provided both more accurate (Figure 9A and Figure 14) and more precise model estimates (shown for two separate models, DTI and ball & sticks – Figure 9B and Figure 11C).

zOverall, the trends identified here extend and complement previous findings (e.g., (Fadnavis et al., 2020; Kay, 2022; Moeller et al., 2021a; Sotiropoulos et al., 2013c; Veraart et al., 2016b)). Yet, the approach followed in this work allowed us to identify differences with respect to previous studies and to define the sources of these differences. For the first time, we defined a gold standard using in-vivo complex averaging that we could use as an ultimate performance reference for denoising approaches. This is not a straightforward task, particularly for high-resolution data (dataset C). Multiple in-vivo repeats of long scans require distortion and motion correction of complex data prior to averaging (as the total scan time for all repeats was ∼1.5 hours), which we addressed here. To ensure that our results arise from differences in the performance of denoising algorithms rather than potential interactions between denoising and pre-processing methods, we intentionally kept parts of the pre-processing pipeline fixed (for instance same susceptibility-induced off-resonance fieldmap was used for a given dataset across all approaches).

With respect to differences between NORDIC and |MPPCA| (Moeller et al., 2021a), we showed that a main source is linked to operating in the complex vs magnitude domain and the ability to deal with the noise floor. Even if the performance of NORDIC and MPPCA* (that differ mainly in the variance normalisation step that NORDIC uses) was not identical, it was considerably more similar than comparing against |MPPCA|. Furthermore, we found detectable spatial resolution penalties in all patch-based methods considered here (Figure 7), which are consistent with similar findings for patch-based denoising of functional MRI data (Vizioli et al., 2021). For the first time, we performed a comprehensive evaluation of different methods against different patch sizes and we found similar penalties and similar trends for all the datasets: the larger the patch-size, the smaller penalty in spatial resolution for PCA-based methods. In theory, larger patch sizes are beneficial for meeting the asymptotic conditions of the Marchenko-Pastur law. However, for too large patches information can become too heterogeneous, redundancy decreases, and this yields to higher deviations from the low-rank assumption, making the determination of the threshold for separating noise from signal more difficult (Moeller et al., 2021a). This can lead to reduced noise level estimates (and removal) and can indirectly also reduce the level of induced covariance. This is an aspect that would need further exploration, taking as well into consideration other factors such as overlapping of the patches. Nevertheless, we found that when an optimal patch size is used a considerable gain in SNR/CNR can be achieved without introducing significant spatial resolution penalties (less than 10% for MPPCA and NORDIC compared to RAW data resolution) even when operating at very low SNR regimes. However, such optimal patch size may not necessarily be the default chosen for specific implementations (Figure 7).

We further explored whether a range of denoising methods are well-conditioned, inducing a larger benefit for noisier data and converging at the high-SNR limit (Figure 10). We found that none of the methods we tested were detrimental to any subsequent processing steps across all datasets and all of them converged towards the behaviour of undenoised data for dataset A (mid to high SNR). We however found that statistical assumptions made for the signal are better fulfilled when denoising in the complex domain, which therefore yielded more accurate DTI estimates against the gold standard complex average.

Gains in signal quality obtained by denoising in the complex-domain can open opportunities for new applications. For instance, we have observed how denoising can turn barely usable data at sub-mm resolution (very low SNR) into data that high-level analyses can run on, like tractography. Denoising can be potentially applied also to reduce the scan time needed to achieve a certain SNR level, either by reducing the need for averages or the need for many dMRI volumes. We found that for an ultra-high spatial resolution dataset, 2/3 of the directions suffice for a number of denoising approaches to converge towards a gold standard obtained using all directions with four repeats; and that 1/3 of the original directions after denoising converge closer to that gold standard than the full raw dataset. This advantage can be applied for both shortening clinical/research scans or pushing the boundaries in SNR-starved applications (e.g., post-mortem imaging).

A valid question is how to treat already acquired or legacy datasets that do not have complex data available. Our results suggest that for relatively good SNR data (e.g., UK Biobank style, such as dataset A), magnitude-based denoising is very comparable to complex-based denoising in terms of secondary-level analysis (e.g., fibre modelling or tractography) as biases from noise floor are small. At the same time, none of them is considerably different to using the raw data directly without denoising, while denoising can induce some spatial resolution penalty. For lower SNR data (e.g., HCP-style, such as dataset B and C), one needs to be fully aware that magnitude denoising preserves noise floor much more than complex domain denoising. And that can bias model estimates such as FA and particularly MD (Fig. 13), while it has been shown that uncertainty in fiber orientations will be inflated as well (Sotiropoulos et al MRM 2013).

### 4.1. The challenge of defining an in-vivo gold standard to compare against

An important secondary outcome from our analyses is highlighting the challenge of defining an appropriate in-vivo gold standard to compare and validate denoising performance against. In the context of recovering SNR, a ground-truth can be defined by multiple averages of the same scan, so that the noise-induced random variability gets averaged out while the signal information remains. This can work under the assumption of no major motion effects and that a set of measurements corrupted by thermal noise will converge to the true signal across repeated acquisitions because they are governed by additive, zero-mean, symmetric noise, i.e. measurements can have high variance but are unbiased. However, magnitude images do not necessarily follow these properties and suffer from elevated noise-floor effects, particularly at low SNR. Averaging data with a noise floor will result into an average with a noise floor (Dietrich et al., 2001; Eichner et al., 2015; Jones and Basser, 2004; Tax et al., 2021). Comparing denoised data against references that contain such noise floor can lead to erroneous validation. A clear example is the results of Fig.13, where the convergence of different methods towards the reference changes completely when the magnitude average is used as a reference (see Suppl. Figure 9), compared to using the complex average as a reference. It is worth pointing out that the mean±std value of MD in the ventricles at b=1000 s/mm^2^ is 2.14±0.2 ×10^−3^ vs 2.76±0.3 ×10^−3^ mm^2^/s for |AVG| and AVG* respectively (with an anticipated self-diffusion coefficient of water in a barrier-free medium at body temperature of ∼3×10^-3^ mm^2^/s); highlighting the level of bias that noise floor can produce in the |AVG| signal. Similarly, the interpretation of how the denoised angular power spectrum looks like (Figure 3) can be challenging without considering a complex average to highlight that power changes in low orders are due to suppression of noise floor.

Having an appropriate gold standard allows unbiased and direct performance metrics and sets the foundations for having a scoring system. This is of crucial importance for comparing denoising approaches given the plethora of methods proposed in the last years (see Suppl. Figure 1). There have been some previous attempts to define a scoring system. For instance, (Chow and Rajagopal, 2017) adapted the blind/reference-less image spatial quality evaluator (BRISQUE) (Mittal et al., 2012) for structural MRI, based on image features such as luminance. In dMRI, (Fadnavis et al., 2022b) recently proposed the Noise Uncertainty Quantification (NUQ) metric, where the Bayesian framework proposed by (Sjölund et al., 2018) is used to develop an uncertainty score for parameters estimates; the lower the noise in the data, the lower the uncertainty expected. Thus, the NUQ can be used to indirectly compare denoising methods. However, as we saw before, reduction of variance is not the only aspect that one should consider to characterise denoising performance. Here we are providing a comprehensive framework that highlights aspects of performance that have received less attention but are still crucial for characterising signal properties after denoising and their implications for downstream analysis.

### 4.2. Denoising as filtering or deterministic prediction?

Denoising is a filtering operation, where an algorithm aims to identify components of the measured signal that represent noise and separate those (filter them out) from the true signal. PCA-based approaches, such as MPPCA and NORDIC, are designed to inherently follow this concept, following in spirit similar concepts in other modalities, such as ICA-based denoising of resting-state functional MRI data (Salimi-Khorshidi et al., 2014) (where noise components are explicitly regressed out from the signal).

A slightly alternative view is that model fitting can be considered a denoising strategy on its own and one can denoise based on a forward (non-)parametric prediction, e.g. (Xiang T. et al., 2023; Jurek et al., 2023a; Tian et al., 2021; Fadnavis et al., 2020). Following this approach, the measured signal informs a prediction maker, which generates a deterministic sample from what the data for a diffusion-weighted volume should look like given all the other volumes. This has the obvious benefit that no patches are used anymore (contrary e.g., to MPPCA or NORDIC) and, as we saw in Figure 7, this could have the potential to result in maximum SNR gains with minimal resolution penalties. At the same time, such approaches have the drawback that they replace the whole data with a single deterministic prediction. By definition, they will be noise-free (as all deterministic predictions are) and hence will have inflated precision. This can have implications on subsequent processing steps (e.g., biophysical model fitting routines or image pre-processing) that assume certain statistical properties for the signal. In addition, it raises the question of which deterministic prediction is the most appropriate. For instance, using a simple linear regression, any microstructural model (such as DTI or DKI), spherical harmonics or any other basis set, Gaussian Processes are all valid options, which could bias towards the specific model used. Hence, replacing data altogether with predictions is more invasive than a signal-filtering operation and caution is needed. Even cutting-edge deep-learning approaches, such as generative neural networks, if optimised for predictive performance (as commonly done) rather than signal fidelity, can induce undesired biases (Shmueli, 2010). In this work, we have not considered any of these predictive approaches, but we have proposed both data and evaluation criteria that would be applicable and informative for such methods as well.

### 4.3. Limitations and future work

We have used an exemplar set of denoising algorithms, so specific findings can differ when applying different approaches. Nevertheless, the main trends are expected to be the same and the proposed considerations are not specific to any algorithm or implementation. Hence, different flavours or whether hybrid approaches that combine two or more methods are advantageous (e.g., as suggested in (Mishro et al., 2021)) can be explored following the criteria introduced here.

The denoising methods used were "off-the-shelf" (default) implementations. We have seen considerable variations in the results by just changing one of the hyper-parameters (e.g., the patch size); fine-tuning of parameters and optimising approaches can be achieved using the framework we presented in this study. We however expect main trends to be preserved regardless of such fine-tuning.

A number of interesting questions regarding denoising applications remain open and we list here some of them as potential future work using the framework proposed. For instance, the interaction of denoising with other pre-processing steps can be evaluated (especially those that increase the rank of the non-thermal noise matrix and affect PCA-based methods), as explored with phase-correction (Jurek et al., 2023b; Liu et al., 2022; Cole et al. 2021; Pizzolato et al., 2020; Cordero-Grande et al., 2019) or motion correction previously (Cieslak et al., 2022; Moeller et al., 2021a; Schilling et al., 2021). In addition, it would be interesting to explore whether signal transformations aiming to reduce the noise-floor bias "post-denoising" (e.g., Koay’s method of moments (Koay and Basser, 2006; Koay et al., 2009), the Variance Stabilization Transform (Ma et al., 2020b)), including the noise-floor as part of the model fitting (Jbabdi et al., 2012)) are as advantageous as performing denoising in the complex domain, as this has been generally explored only for magnitude-based denoising (Hutchinson et al., 2017). We anticipate that even if mitigation of noise floor bias effects is possible through post-denoising approaches, they have the obvious disadvantage that any signal rectification caused by noise (e.g. Figure 5) is irreversible. Hence, solving the problem would always be preferable to mitigating its effects. In terms of scan-rescan reproducibility, thermal noise can be a major driver of variability in dMRI experiments (Schilling et al., 2021). We therefore anticipate this variability to be reduced after denoising although further exploration is needed. Finally, exploring the performance of denoising for non-conventional dMRI sequences, such as Double-Diffusion Encoding (Jespersen et al., 2013), q-space trajectory (Westin et al., 2016) or Oscillatory Gradient Spin Echo (Does et al., 2003) would be interesting, as the level of signal redundancy for these could be very different to conventional PGSE sequences.

We hope this work paves the way for future extensions where the proposed criteria can be formalised and integrated into an automated scoring system. This would help in development and optimisation of new methods and performance comparison across methods. Given the lack of ground-truth for denoising (even if complex space powder-averaging gives a good approximation to it), a potential line of research is the application of unsupervised machine learning to learn no-reference scores (e.g., (Chow and Paramesran, 2016; Lin et al., 2020; Stępień et al., 2021)), such as the Modified-BRISQUE for MRI (Chow and Rajagopal, 2017). This type of scores can offer a more sophisticated alternative to SNR or CNR, where aspects such as spatial resolution penalties or noise-floor could be included.

## ACKNOWLEDGEMENTS

JPMP and SNS acknowledge funding from the European Research Council (ERC Consolidator - 101000969 to SNS). JPMP was also partially supported by a University of Nottingham PhD studentship from the Precision Imaging Beacon of Excellence. JA is supported by the Wellcome Trust [203139/Z/16/Z]. SM, KU and EY are supported by the National Institutes of Health (NIH grants P41EB027061, U01EB025144), RF1 MH116978).

## CONFLICTS OF INTEREST

NORDIC and NIFTI_NORDIC are copyrighted by Regents of the University of Minnesota and covered by issued US patent #10,768,260, and S.M. has a relationship with this patent. S.M did not evaluate or analyse any of the data in this work. No other authors have competing interests to declare.

## DATA AND CODE AVAILABILITY

Data are available as an OpenNeuro repository. Preprocessing was done using standard FSL tools (FSL v 6.0.6, CUDA 10.2), used as in the publicly released HCP pipelines (Glasser et al 2013, Neuroimage). We specifically used an in-house version of this pipeline: https://zenodo.org/record/8059550 <u>(v. 1.6.3)</u>.

For denoising, DIPY v1.7 (https://dipy.org/documentation/1.4.1./examples_built/denoise_nlmeans/) was used for NLM, mrtrix3 (https://mrtrix.readthedocs.io/en/dev/reference/commands/dwidenoise.html) for MPPCA, and the released implementation (https://github.com/SteenMoeller/NORDIC_Raw) for NORDIC.

We also share a Jupyter notebook for reproducing the results provided in the manuscript (python 3.9), available on https://github.com/SPMIC-UoN/EDDEN. Code for phase processing and projection of complex data to real space is available in the same repository.

## Notes

### Competing Interest Statement

The authors have declared no competing interest.

### Summary of Updates

Added comments from reviewers and access to code repositories.

