## Supplementary Material for "Denoising Diffusion MRI: Considerations and implications for analysis"

#### *Supplementary Text*

##### *Phase-correction for rotating complex-valued images to the real channel*

The processing of the complex data prior to denoising is applied with the intent to remove most large phase-fluctuations and to have the majority of signal in the real channel and the imaginary channel being largely (but not exclusively) noise. The processing is performed in 2 steps for the static and dynamic phase changes, respectively. First, a slice-specific volume independent correction is applied in which the average unfiltered phase is removed; this removes static effects related to B0, sensitivity profiles and partial Fourier sampling. Secondly, a slice-specific volume dependent correction is applied in which a smooth phase is estimated to largely change the acquired complex data; this removes the motion induced phase variations during the diffusion gradients.

Code for this processing is available on:
https://github.com/SPMIC-UoN/EDDEN/blob/main/NIFTI_COMP_to_REAL.m

##### *Software and denoising settings*

| ***Software*** | ***Version*** | ***Command*** | ***Settings*** |
|---|---|---|---|
| ***NLM (DIPY)***<br><br>*https://dipy.org/* | *1.7.0* | *dipy_denoise_nlmeans* | *Default: Gaussian kernel, patch_radius=1 (patch_size = 2*patch_radius+1), block_radius=5* |
| ***MPPCA (MrTrix)***<br>*https://www.mrtrix.org/* | *3.0.4* | *dwidenoise* | *Default: extent=P†, estimator=Exp2* |
| ***NORDIC***<br>*https://github.com/ SteenMoeller/NORDIC_Raw* | - | *NIFTI_NORDIC.m* | *Default: ARG.temporal_phase=1, ARG.phase_filter_width=3, ARG.kernel_size_PCA=[P,P,P]* *§* |

**Suppl. Table 1:** Software and default denoising implementations use*d. For NLM, experiments with the Rician kernel were tried, but these produced excessive intensity suppression in dataset C (where noise floor effects are maximal); this behaviour has been previously reported (https://xgrg.github.io/denoising-DWI/).*

*†For MPPCA, the patch size P is selected automatically in dwidenoise such as P is the smallest isotropic patch size such that the number of contained voxels $P^3$ exceeds the number of DWI volumes N in the data. E.g. P=5 for a 5x5x5 patch for data with N<=125 DWI volumes. This parameter is modified to explore different patch sizes (Fig. 7 and 8).*

*§For NORDIC the patch size P is by default set such that the ratio between the number of voxels in the patch and the number of volumes N is $P^3$/N>=11. E.g., for Dataset C with N=199 directions, P is automatically set to 13. This parameter is modified to explore different patch sizes (Fig. 7 and 8).*

### *Supplementary Figures*

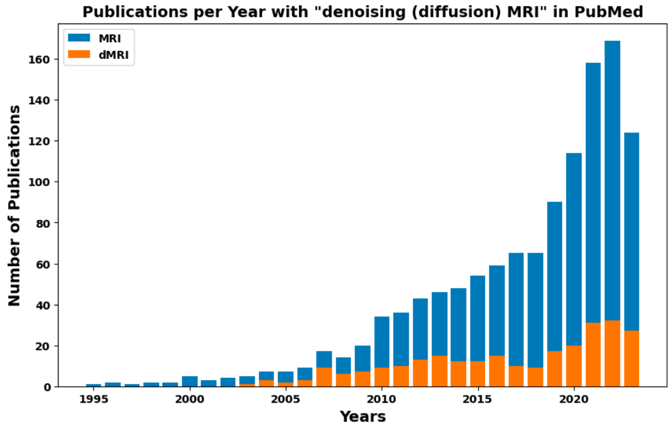


**Suppl. Figure 1.** *Number of publications per year in PubMed containing in the title the words "Denoising MRI" (in blue) or "Denoising diffusion MRI" (in orange). It is updated until October 2023.*

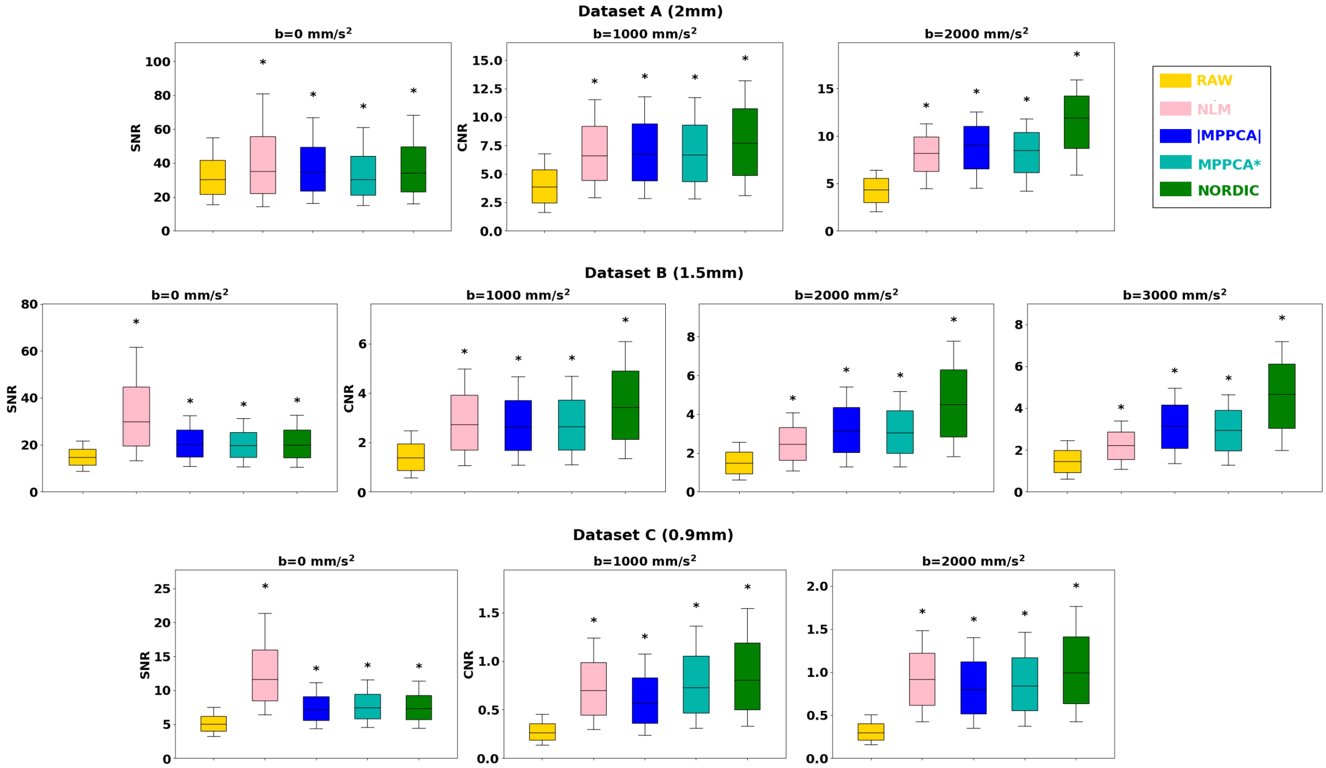


**Suppl. Figure 2.** *Vowel-wise Signal-to-Noise Ratio (SNR) and Angular Contrast-to-Noise Ratio (CNR) in raw and denoised data. Boxplots correspond to the distribution of these metrics in white matter. Top row: Dataset A (2mm), Middle row: Dataset B (1.5mm), Bottom row: Dataset C (0.9mm). Asterisk indicates statistical significance of pairwise Wilcoxon rank-sum tests between raw vs each denoised approach, Bonferroni corrected for multiple comparisons.*

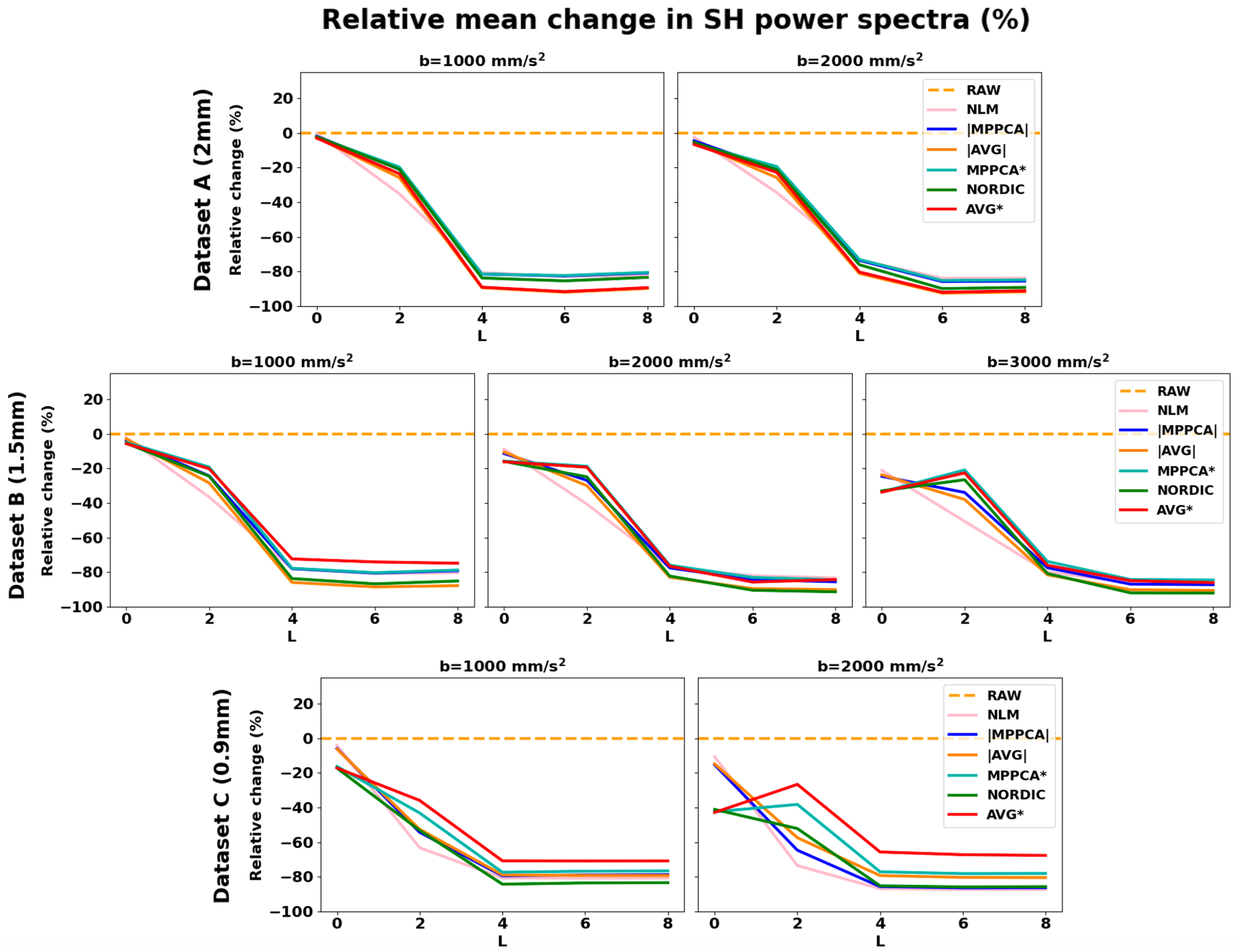


**Suppl. Figure 3.** *Power spectra of Spherical Harmonics. Mean relative change (%) in the gray matter spherical harmonics (SH) power after denoising, with respect to the corresponding power for the raw data. Results for even harmonic orders L=0–8 are presented. Top row: Dataset B (1.5mm), Bottom row: Dataset C (0.9mm)*

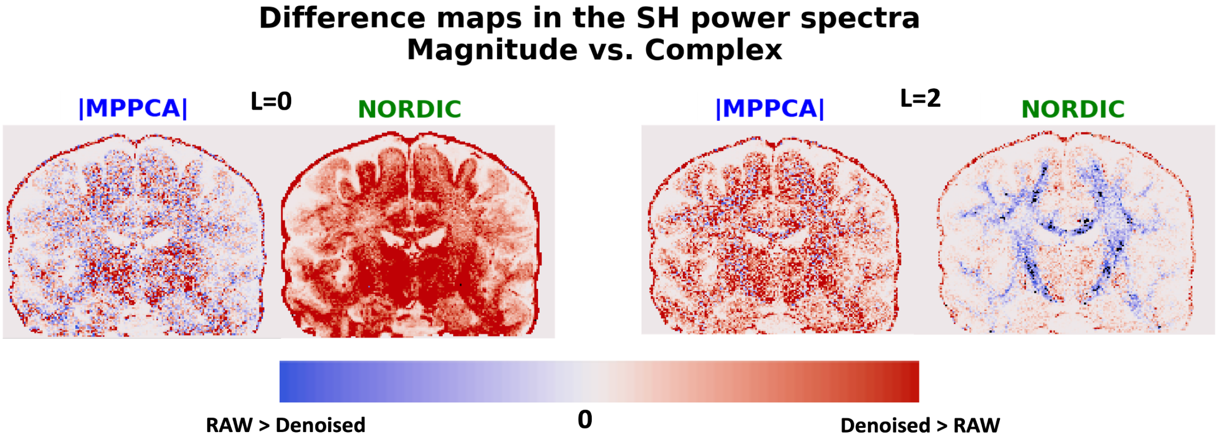


**Suppl. Figure 4.** *Exemplar maps of the differences in the SH power spectra in magnitude- and complex-denoising at orders L=0 and L=2 (Dataset C, b=2000 s/mm²). Reduction of L=0 power (flat signal) and increase in L=2 power (unimodal anisotropic signal) with complex-domain approaches can be associated with the signal recovered by pushing lower the noise-floor.*

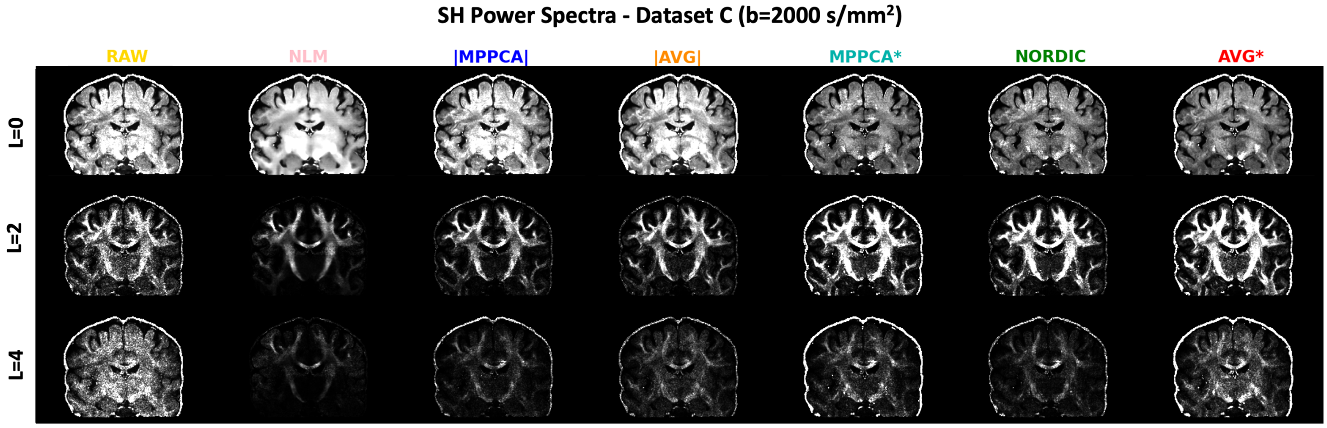


**Suppl. Figure 5**. *Exemplar maps of the white matter power spectra of even harmonics orders L=0–4 Spherical Harmonics for Dataset C (0.9mm) in the highest b-shell (b=2000 s/mm²). For each row the same minimum and maximum intensity limits have been used for visualisation, so brightness of intensity corresponds to absolute power for each case (i.e., denoised data reduces power at higher orders L).*

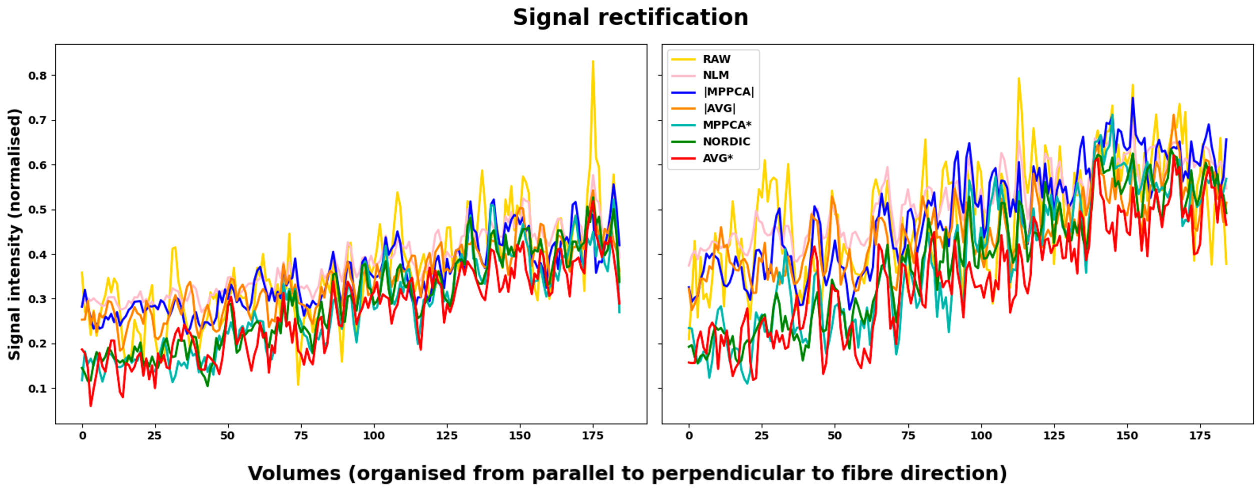


**Suppl. Figure 6.** *Signal intensity values in two example high FA voxels (FA>0.8) from the Corpus Callosum. Signal for different diffusion-sensitising volumes has been ordered so that gradient orientations are from parallel to perpendicular to the primary fibre orientation (DTI* $v_1$*) of AVG*, i.e. approximately lowest to highest signal intensity. The effect of signal rectification and loss of signal dynamic range in the magnitude domain are apparent and these are preserved with all magnitude-based approaches. For better visualisation, a moving average using three neighbouring points has been used in these plots.*

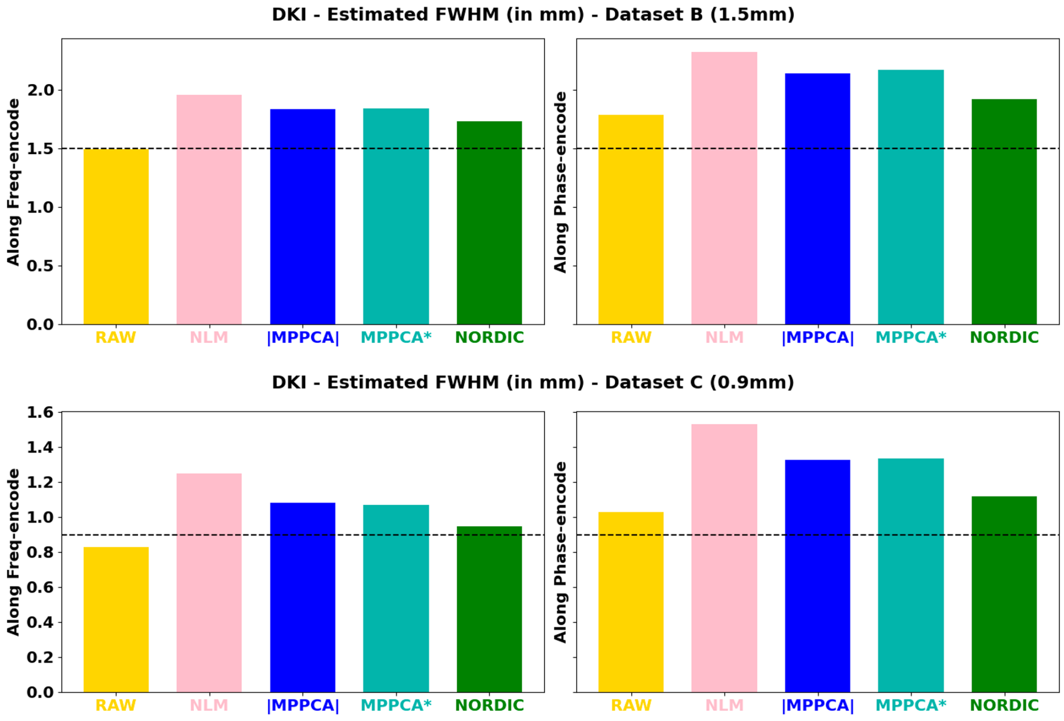


**Suppl. Figure 7.** *Estimated voxel resolution for the denoised and non-denoised datasets, using DKI residuals of multishell data, along the frequency (acquisitions along x-axis) and encode (acquisitions along y-axis) directions - Top row: Dataset A (2mm); Middle Row: Dataset B (1.5mm); Bottom row: Dataset C (0.9mm). Dashed line: acquisition nominal resolution.*

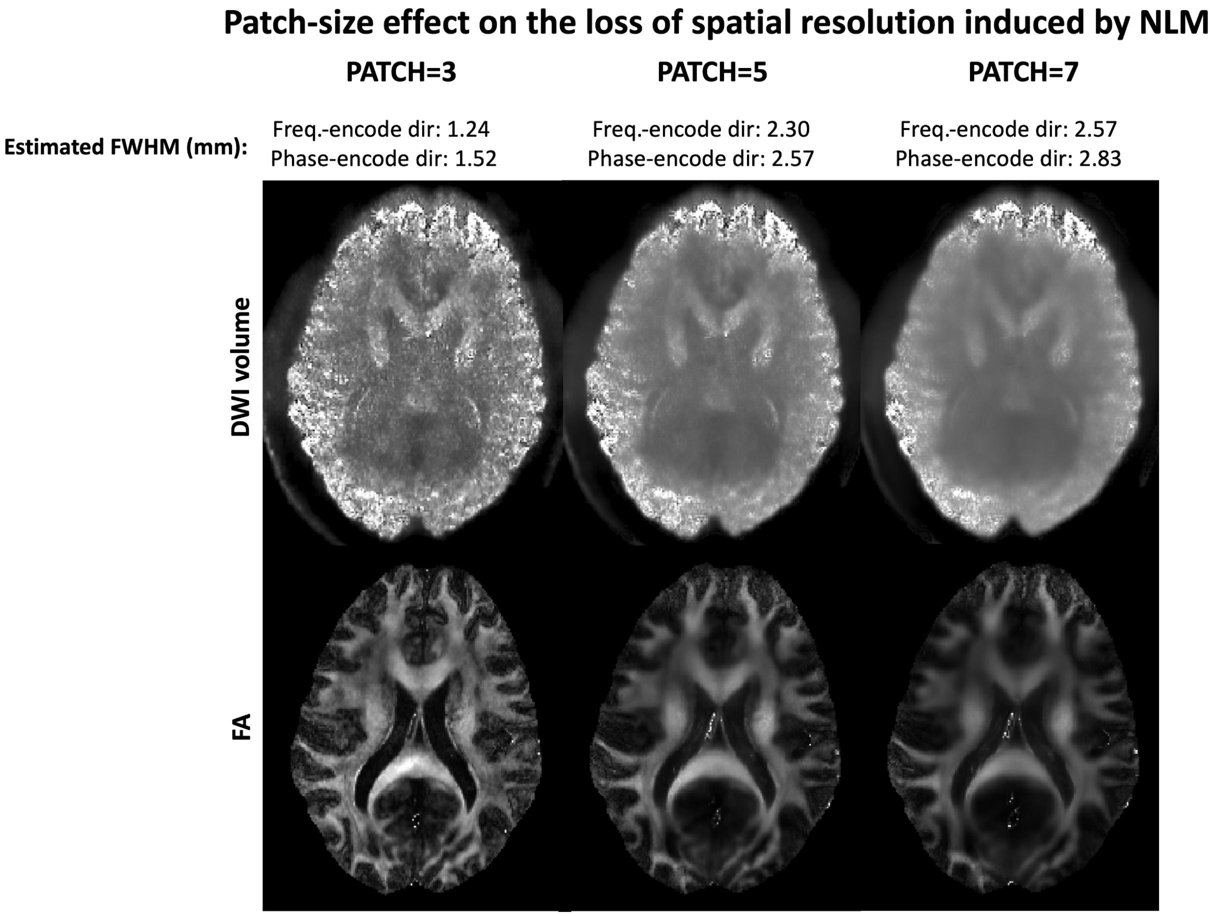


**Suppl. Figure 8.** *Effect of patch-size increase in NLM denoised data (top row) and derived FA maps (bottom row). The spatial smoothing introduced by NLM increases rapidly with the size of the patch, to the point that it is barely usable out of the default patch-size of a 3x3x3 neighbourhood (leftmost column). In all cases we maintained a ratio 1:5 for patch_radius vs block_radius, block size being the area of the image that is searched for similar voxel patches and allows computational feasibility.*

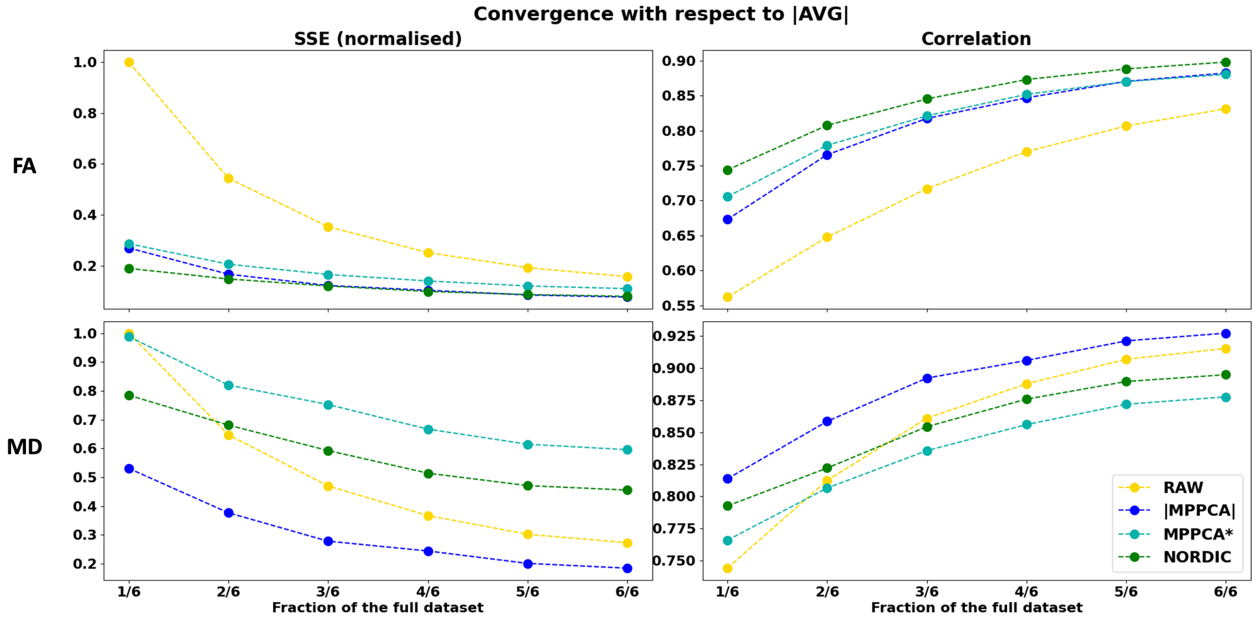


**Suppl. Figure 9.** *Convergence to the magnitude multiple-averages (|AVG|) assessed by the Sum of Squared errors (left column) and Pearson correlations (right column) in DTI model estimates of subsets from Dataset C (0.9mm). Top: Fractional Anisotropy correlations. Bottom: Mean Diffusivity.*
